# Species wide inventory of *Arabidopsis thaliana* organellar variation reveals ample phenotypic variation for photosynthetic performance

**DOI:** 10.1101/2024.07.12.603232

**Authors:** Tom P.J.M. Theeuwen, Raúl Y. Wijfjes, Delfi Dorussen, Aaron W. Lawson, Jorrit Lind, Kaining Jin, Janhenk Boekeloo, Dillian Tijink, David Hall, Corrie Hanhart, Frank F.M. Becker, Fred A. van Eeuwijk, David M. Kramer, Erik Wijnker, Jeremy Harbinson, Maarten Koornneef, Mark G.M. Aarts

## Abstract

Efforts to improve photosynthetic performance are increasingly employing natural genetic variation. However, genetic variation in the organellar genomes (plasmotypes) is often disregarded due to the difficulty of studying the plasmotypes and the lack of evidence that this is a worthwhile investment. Here, we systematically phenotyped plasmotype diversity using *Arabidopsis thaliana* as a model species. A reanalysis of whole genome resequencing data of 1,531 representative accessions shows that the genetic diversity amongst the mitochondrial genomes is eight times lower than amongst the chloroplast genomes. Plasmotype diversity of the accessions divides the species into two major phylogenetic clusters, within which highly divergent subclusters are distinguished. We combined plasmotypes from 60 *A. thaliana* accessions with the nuclear genomes (nucleotypes) of four *A. thaliana* accessions to create a panel of 232 novel cytonuclear genotypes (cybrids). The cybrid plants were grown in a range of different light and temperature conditions and phenotyped using high-throughput phenotyping platforms. Analysis of the phenotypes showed that several plasmotypes alone or in interaction with the nucleotypes have significant effects on photosynthesis, and that the effects are highly dependent on the environment. Moreover, we introduce Plasmotype Association Studies (PAS) as a novel method to reveal plasmotypic effects. Within *A. thaliana,* several organellar variants can influence photosynthetic phenotypes, which emphasizes the valuable role this variation has on improving photosynthetic performance. The increasing feasibility of producing cybrids in various species calls for further research into how these phenotypes may support breeding goals in crop species.

**Significance statement:** Photosynthesis is one of the few crop traits that has been largely unaddressed which can contribute to increasing crop yield potential. Exploiting genetic variation within organellar genomes presents a promising, yet untapped resource to improve photosynthesis. However, the extent of organellar variation and its impact on photosynthesis within a species remains largely unknown. Using *Arabidopsis thaliana* as a model species, we revealed highly divergent clusters of organellar variation. We constructed 232 novel combinations of species representative organellar and nuclear genomes, referred to as cybrids. High-throughput phenotyping of these cybrids revealed that organellar variants can substantially impact photosynthesis in different environments. These findings indicate that organellar genomes may be a valuable resource for improving photosynthesis in crops.

## Introduction

Improving photosynthetic performance has become an important goal for increasing crop yield to meet global food demands in the coming decades (1–4). Genetic variation is often overlooked in studies on improving photosynthetic performance (5, 6) due to the limited variation in genes encoding the core photosynthetic machinery (3). Recently, evidence is accumulating that genetic variation can contribute to photosynthetic differences within species (7–18). Even though this research area is rapidly gaining more attention, the contribution of the organellar genomes to photosynthetic variation remains unclear. In plants, this organellar genetic variation is divided between two organelles that each have their own genome, the chloroplast and the mitochondrion. Derived from ancient endosymbiotic ancestors, the genomes of the extant organelles are drastically reshaped by evolution. Thousands of genes were lost or moved to the nuclear genome, while only some genes have been retained in the organellar genomes (19). The chloroplast and mitochondrial genomes of *Arabidopsis thaliana* only harbour 129 and 57 genes, respectively (20, 21). The retention of genes in the organellar genomes may be explained by the need for cross talk between the organellar and nuclear genomes; suppressed mutation rate in organellar genomes; difficulty in importing proteins over the organellar membranes; or the requirement of redox-regulated gene expression (22). The retained genes primarily encode components of the photosynthetic and respiration machinery and essential components needed for organellar gene expression.

Genetic variation in the organellar genomes may be limited as a result of purifying selection and neutrality of variants (23). This would explain the strong conservation of organellar genes across plant species and the observations that mutations in organellar genes often lead to severe, debilitating phenotypes (24, 25). The reduced number of nonsynonymous mutations in comparison to synonymous mutations is further evidence that purifying selection is prominent in organellar genomes (26, 27). The mutation rate of a plant mitochondrial genome is generally regarded to be approximately sixfold lower than that of a nuclear genome, while the chloroplast genome mutation rate is about twofold lower than that of the nuclear genome (28). Over time though, mutations are expected to occur, causing organellar genetic variation that can be selected for in natural environments. An example of organellar adaptation is the mutation in *PsbA*, which encodes the D1 protein of photosystem II (PSII), that confers herbicide tolerance to *A. thaliana* (29). This mutation has been selected for on the British Railway network through application of herbicides (30). Another example is the *RbcL* gene, encoding the Rubisco large subunit, where several amino acid residues are under positive selection in land plants (31). Organellar variation in *Arabidopsis lyrata* is also found to increase fitness in specific environments (32). Systematic analyses of organellar variations within a species are rare, but studies using *A. thaliana* suggest that cytoplasmic variation is widespread (33–36). This shows that organellar variation can contribute to plant adaptation, even though it is slower and more limited than adaptation based on nuclear variation.

To assess the role of organellar genetic variation in photosynthetic performance, a systematic analysis of organellar genotype and phenotype variation at the species level is needed. However, the cytoplasm is uniparentally inherited, which complicates efforts to separate the nuclear and cytoplasmic origins of phenotypic variation. Even though reciprocal hybrids and segregating populations can give a good indication of cytoplasmic effects, maternal effects and genomic imprinting are known to play a significant role and could be interpreted as false-positive cytoplasmic effects (37–39). To exclude maternal effects and genomic imprinting, recurrent backcrossing can be used. This approach allows the production of novel combinations (cybrids) of nuclear genomes (nucleotypes) and organellar genomes (plasmotypes). Analysis of the resulting cybrids has indeed shown that phenotypic variation can be associated with organellar genomes in different plant species (40–42). Backcrossing approaches are particularly useful in plant breeding programs, but they are lengthy and residual nuclear introgressions that can influence phenotypic variation may be remaining. Flood *et al.* (2020) showed that in *A. thaliana* a maternal haploid induction system can be used as an attractive alternative method to create cybrids without a lengthy backcrossing procedure (43). Such maternal haploid induction system is based on a *GFP-tailswap* mutant, where the centromere-specific histon 3 (*CenH3*) gene is replaced by a *GFP*-tagged *CenH3* gene. A *GFP-tailswap* mutant can be used to replace the complete nucleotype within one generation (43, 44). This method can efficiently be used to systematically separate phenotypic variation caused by nucleotype variation from that caused by plasmotype variation.

Flood *et al.* (2020) showed that plasmotypic variation can result in significant differences in photosynthetic performance (43). These so far unknown phenotypic differences were found to be caused either by variation in the plasmotype alone or as a result of an interaction between plasmotype and nucleotype variation. The phenotypic differences were most often observed in dynamic light conditions. This showed that photosynthesis is affected by plasmotype variation, but the seven accessions of the cybrid panel represented a limited fraction of the plasmotype diversity to be found in *A. thaliana*. As *A. thaliana* is native to most of the Eurasian landmass and high-altitude regions of Africa, accessions originating from vastly different environmental conditions are available (35, 45–49). The availability of accessions that potentially represent the global species-wide genetic diversity, makes *A. thaliana* an excellent model system to explore the extent to which plasmotypic variation contributes to overall phenotypic variation, and more specifically photosynthetic variation.

Here, we reanalyse the whole-genome sequencing data of 1,541 publicly available *A. thaliana* accessions to obtain high quality organellar genetic variation. Using this dataset, we reveal how much genetic variation is present within the species and how it is distributed over the organelles. From this dataset, a selection of 60 genetically diverse, species-representative plasmotypes were combined with four distinct and diverse nucleotypes to construct a new panel of 232 cybrids. To characterize and understand the contribution of plasmotype variation to photosynthetic variation we phenotyped this cybrid panel in detail using three different experimental set-ups where they were exposed to a range of light and temperature conditions.

## Results

### Plasmotypic variation analysis

To quantify the plasmotype diversity among *A. thaliana* accessions, we reanalysed available sequencing data of 1,541 accessions collected in Europe, Asia and Africa. Variant calling for the organellar genomes of a subset was performed previously, but quality control filtering steps were applied as tailored to nuclear genome analyses. Such filtering can result in low quality variant data, since organellar genomes have different intrinsic properties, for example higher copy number and heterozygosity. To compare variants amongst accessions without introducing biases from separate variant calling projects, we performed variant calling on all accessions simultaneously, with quality filters separately tailored to the chloroplast and mitochondria. The updated version of the mitochondrial reference genome was used for variant calling (21).

We first filtered out poor-quality raw sequencing data and accessions with a high percentage of heterozygous calls in the chloroplast genome, which are indicative of sample contaminants. This analysis resulted in variant calling data for the organellar genomes of 1,531 *A. thaliana* accessions. The dataset comprises 5,015 variants in the chloroplast genomes and 1,430 variants in the mitochondrial genomes, compared to the Col-0 reference genome. Resultingly, 3.2% of the nucleotides in the chloroplast genome and 0.4% of the nucleotides in the mitochondrial genome showed genetic variation within at least one accession. Therefore the genetic diversity within the chloroplast genomes is eightfold higher compared to the mitochondrial genomes. Specifically, in chloroplast genes, the ratio between predicted non-synonymous mutations versus synonymous mutations is 0.9, while for mitochondrial genic regions this is 1.8. Thus, we conclude that the genetic diversity in the chloroplasts is higher than that in the mitochondria, but that the chloroplast genes are more conserved than the mitochondrial genes.

Next, we studied the plasmotype diversity between accessions to reveal phylogenetic relationships. The mitochondrial genome is known to be more dynamic than the chloroplast genome (50, 51). This results in extensive interspecies modifications in the mitochondrial genome (52), and therefore may not represent the true phylogenetic origin (33, 53, 54). Therefore, we focussed on the chloroplast genomes to reconstruct the phylogeny of *A. thaliana*. We found that most accessions were unique in that 1,495 haplotypes could be identified among the 1,531 accessions. Principal component (PC) analysis of the chloroplast variation revealed three distinct main clusters diverging from one central point, but the fraction of variation explained by PC1 (2.1%) and PC2 (1.5%) is low (Figure 1A). Rooting the neighbour joining tree based on the chloroplast genome with *A. lyrata*, *Arabidopsis halleri*, *Arabidopsis carpatica* and *Capsella rubella* shows that *A. thaliana* accessions have diverged into two main clusters (Figure 1B). With a cophenetic correlation coefficient of 0.65, the neighbour joining trees of the chloroplast and mitochondrial genomes have substantial overlap (Supplementary Figures 1 and 2). On the other hand, the cophenetic correlation coefficient between the neighbour joining trees for the chloroplast and nuclear genomes is relatively low at 0.26 (Supplementary Figures 1 and 3). The deep branch lengths in the chloroplast neighbour joining tree and the low fraction explained by the main PCs indicate there is substantial genetic variation between different clusters.

**Figure 1.**
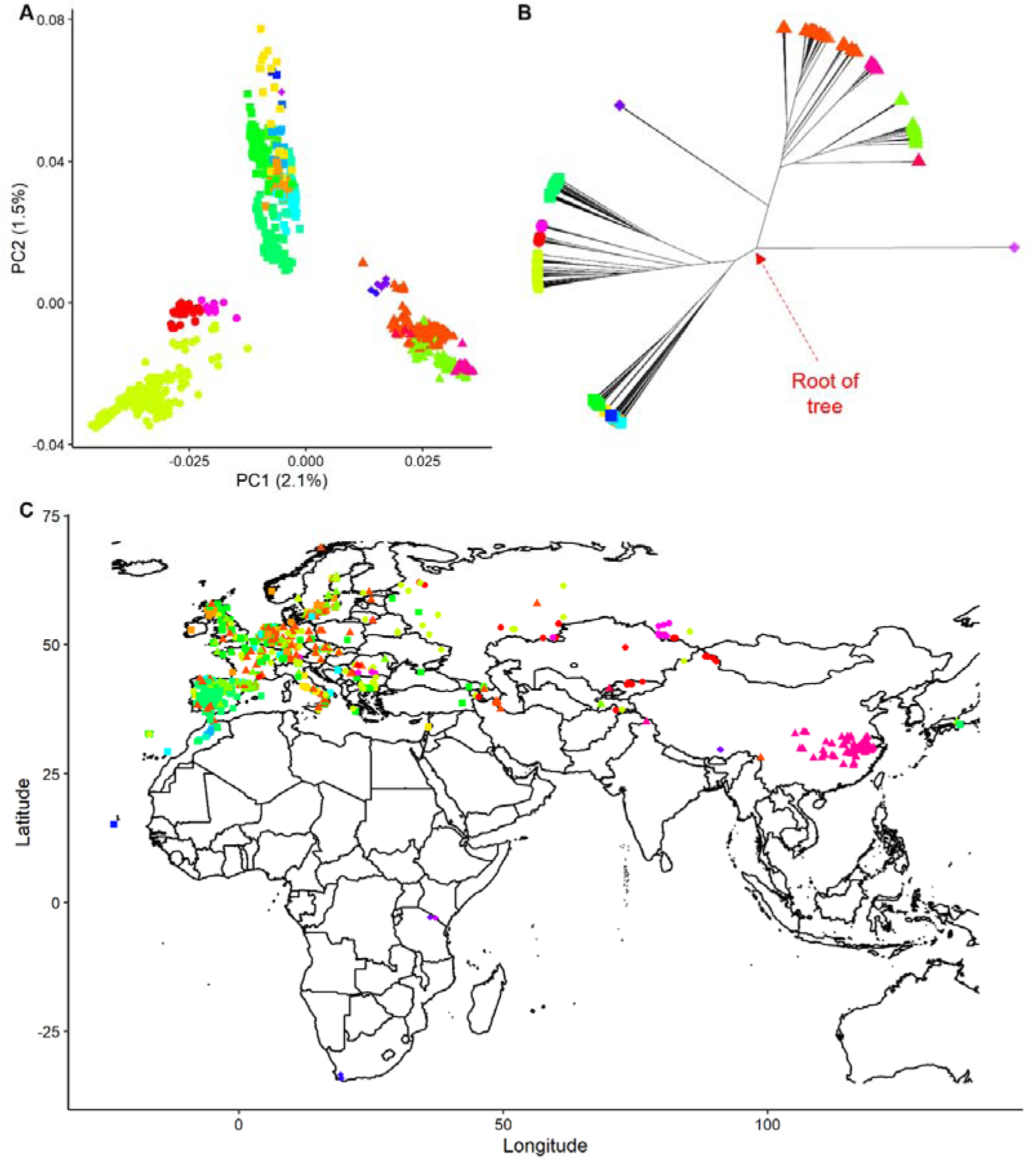
Phylogenetic analyses of organellar diversity in A. thaliana based on chloroplast genome variation. A) The first two principal components (PCs) showing the major components that separate the organelle diversity into three main branches. Colours are based on hierarchical clustering on the basis of a neighbour joining tree with k=20. Accessions within each of the three branches are depicted with a triangle, circle or square, with the exception of accessions close to the root of the neighbour joining tree. The same colour codes and symbols are used in panels B and C. For additional PCs see Supplementary Figure 5. B) Neighbour joining tree displaying the genetic relationship between accessions, based on chloroplast genetic diversity. The red arrow indicates the root of the tree, where the four related species are connected. For clarity these are left out. C) Accessions with colours and symbols as defined in panel A and B mapped to the geographic positions where they have been collected. Accessions sampled in North-America are not shown as these are found to have originated on the European continent (45).

Much of the observed organellar variation could be the attributed to population structure and historical species expansion. To define different subclusters we used the elbow method and arbitrarily chose k = 20 (Supplementary Figure 4). Organellar subclusters could arise in genetic isolation without geographic isolation due to the inability to recombine amongst accessions. Plotting the accessions onto their geographic locations shows how accessions from West Asia and Europe are largely a mixture of the most common organellar clusters observed within *A. thaliana* (Figure 1C). In other geographic locations only specific organellar subclusters occur: the Cape Verde islands and Madeira each represent their own subcluster, and also the Yangtze River basin, the Northern and Southern Altai mountains, Uzbekistan, the Iberian Peninsula, Tanzania, South-Africa and several regions within Morocco are home to unique accessions from one subcluster. This shows that in some parts of the native range, organellar subclusters are specific to one geographic location and absent in other regions. The geographic isolation of clusters may be due to neutral mutation accumulation, but it can also result in genetic adaptation.

### Novel cybrid panel construction

We constructed a novel cybrid panel to generate a representative overview of the impact of organellar variation on plant phenotypes beyond the previous cybrid panel we examined (43). We selected 53 accessions from 18 of the 20 chloroplastic subclusters we defined, from which seeds were available. The two subclusters not represented contain accessions from South Africa and Tanzania and are only known from herbarium material (46). Amongst the 53 accessions is the Staro-2 accession, which represents an outgroup based on the mitochondrial neighbour joining tree (Supplementary Figures 1 and 2). We added seven additional accessions for a total of 60, including three accessions from Africa (i.e. ET2, Tanz-2 and Toufl) for which seeds were available but which had not been sequenced previously. We combined the plasmotypes of all 60 accessions with four divergent nuclear accessions (i.e., Bur-0, Col-0, Tanz-1 and Cvi-0; Supplementary Figure 3). We expected that these will result in many novel nucleotype-plasmotype interactions. In the process of cybrid production we observed substantial variation in the percentage of haploids recovered from the total number of germinating seeds with fractions ranging between 20.1% haploids for the Bur-0 nucleotype to 51.8% haploids for the Col-0 nucleotype. This suggests there is genetic variation for the haploidization based on the *GFP-tailswap* mutant system (55) due to the loss of maternal chromosomes during postzygotic mitotic division. The resulting 240 cybrids were whole-genome-sequenced to verify their genotypes, and 232 cybrids were found to have the expected genotype (Supplementary Figure 6). These 232 cybrids were subsequently used to assess the plasmotypic contribution to photosynthetic variation.

### Cybrids in dynamic environments

Previous work showed that plasmotypes, and the interaction between plasmotypes and nucleotypes, primarily contributed to the overall phenotypic variation when grown under high and dynamic light conditions (43). Therefore, we exposed the cybrids to a range of different conditions including: steady-state light intensities, dynamic light conditions, low temperatures and combinations of these. During these treatments, we phenotyped the photosynthetic response using two different high-throughput chlorophyll fluorescence phenotyping systems (Figure 2 and Supplementary Figures 7 and 8). We phenotyped for F_v_/F_m_, NPQ, NPQ_(t)_, Φ_PSII_, Φ_NPQ_, Φ_NO_, *q*_E_, *q*_E(t)_, *q*_I_, *q*_I(t)_, *q*_L_ and projected leaf area with the Dynamic Environment Phenotyping Instrument (DEPI) and Φ_PSII_ and projected leaf area in the Phenovator system (56, 57). Broad sense heritability (H^2^) was used to estimate how much of the phenotypic variation was explained by the total genetic component. The contribution of the nucleotype, plasmotype and nucleotype-plasmotype interaction to phenotypic variation was calculated as a fraction of the total H^2^. The Ely plasmotype was excluded from H^2^ analysis, because the large-effect *PsbA* mutation in the Ely plasmotype influences the H^2^ to such an extent that it masks observations for the rest of the panel. To determine whether these phenotyping systems allowed us to assess genotype-environment interactions, we first analysed overall H^2^. NPQ was found to have an average H^2^ of 9.2% under steady-state low light, while under fluctuating light this increased to 15% (Figure 2B and Supplementary Table 1). The H^2^ increased further to 34.6% when combining steady-state low light with a low temperature (12 °C). Fluctuating light in combination with a low temperature resulted in H^2^ of 27.5%, with outliers during the high light fluctuations of 61.1%. This shows that our phenotyping setup acts as a reliable platform for observing genotype-environment interactions.

**Figure 2.**
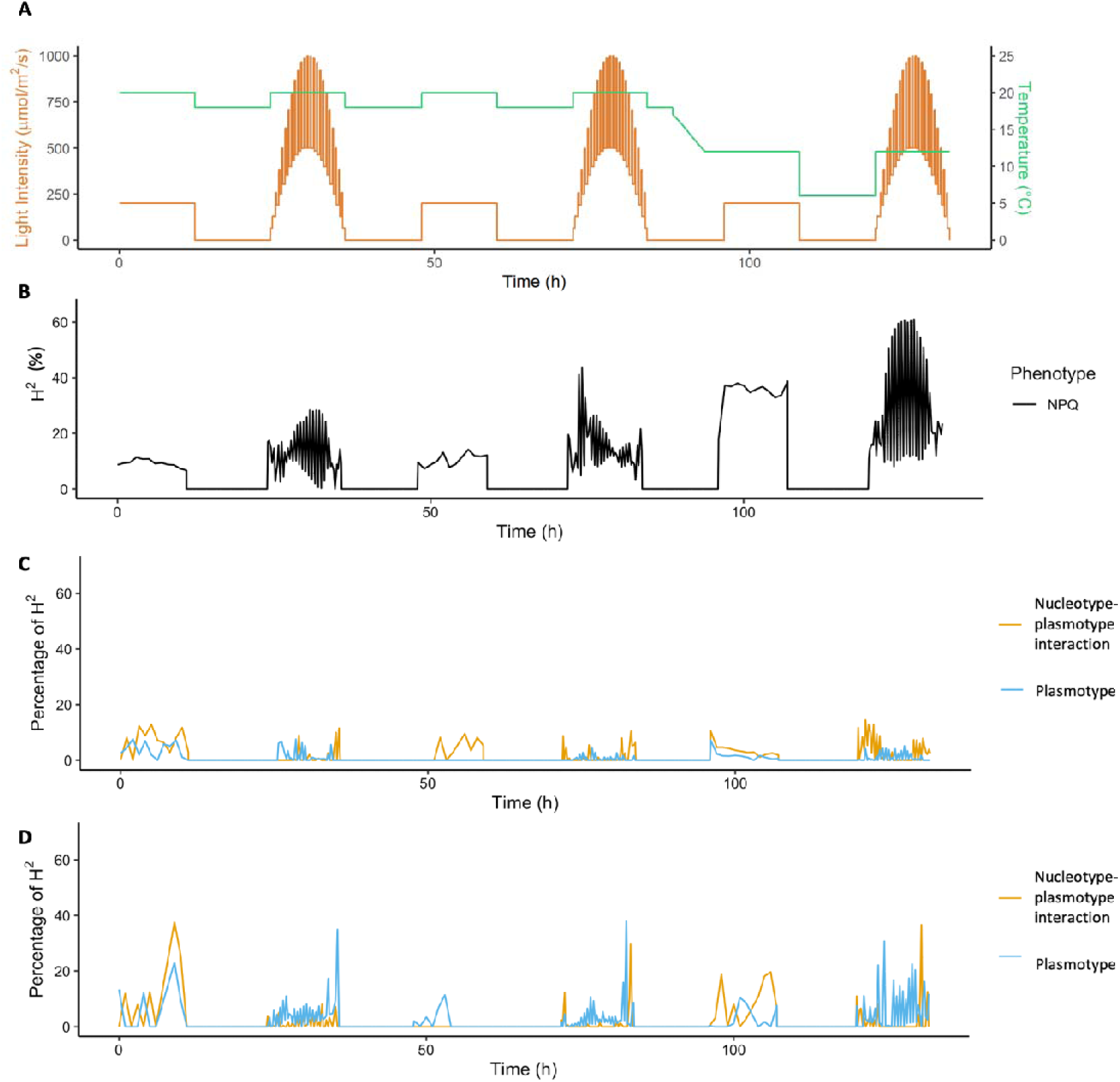
Overview of overall broad sense heritability, and the fractions explained by the plasmotype and nucleotype-plasmotype interaction components of non-photochemical quenching (NPQ) in response to different environmental conditions in the DEPI system. A) Environmental conditions to which the plants were exposed after being grown under steady-state light conditions of 200 µmol m^-2^ s^-1^, with t=0 being the start of the photoperiod 21 days after sowing. B) Broad sense heritability (H^2^; shown as percentage) of NPQ in response to the different environmental conditions, as shown in panel A. C) Percentage H^2^ for NPQ in response to the different environmental conditions for the additive plasmotype effect (blue) and nucleotype-plasmotype interaction effect (yellow). D) The same as panel C, but for the percentage H^2^ for q_E_ in response to the different environmental conditions.

The average H^2^ was 31.6% across all environmental conditions, for a total of 1,986 phenotypes in the DEPI experiment. The plasmotype and nucleotype-plasmotype interaction components explained 1.3% and 1.9%, respectively (Supplementary Figure 7). Regarding NPQ, the fraction of H^2^ for the additive plasmotype accounts for on average 1.1%, and the fraction H^2^ for the nucleotype-plasmotype interaction accounts for 2.5% (Figure 2C). For some phenotypes, under certain conditions, these components account for substantially more, even though the average H^2^ is low. This was most pronounced for the rapid-relaxing component of NPQ, *q*_E_, for which the additive plasmotype effect explains on average 4.7%, and the nucleotype-plasmotype interaction explains 2.5% (Figure 2D). However, up to 36.6% of the total H^2^ for *q*_E_ is explained by the nucleotype-plasmotype interaction under fluctuating light conditions in combination with a low temperature (Figure 2D). For the additive plasmotype effect this was up to 37.9% of the total H^2^ for *q*_E_, at the end of the fluctuating light days (Figure 2D). This shows that plasmotype variation, either via additive effects or via an interaction with the nucleotype, on average accounts for relatively little of the H^2^ in comparison to the nucleotype. However, under specific conditions, the plasmotype plays a substantial role, and therefore can influence overall photosynthetic performance.

The DEPI experiment allowed us to stress the cybrids with high and fluctuating light conditions in combination with cold temperatures at the end of the growing period. However, the impact of dynamic environmental conditions over the whole growing period remained unknown. We grew the entire cybrid panel in semi-protected conditions during the spring of 2020 and 2021 in Wageningen (The Netherlands, 51°59’20.0”N 5°39’43.2”E). In such conditions, the cybrids were constantly exposed to a dynamic environment. Analysis of the data showed that the resulting 57 photosynthetic and morphological phenotypes had an average H^2^ of 16.6% (Supplementary Figure 9), of which the plasmotype explained 2.5% and the nucleotype-plasmotype interaction explained 5.6% (Figure 3D). The plasmotype and nucleotype-plasmotype interaction explained 37.7% of the H^2^ for NPQ (measured as NPQ_(t)_) after a high to low-light transition. This is substantially more than observed for NPQ in the DEPI experiment (Figure 2C), which means that for the same phenotype the contribution of the plasmotype depends heavily on the environment in which plants are grown.

**Figure 3.**
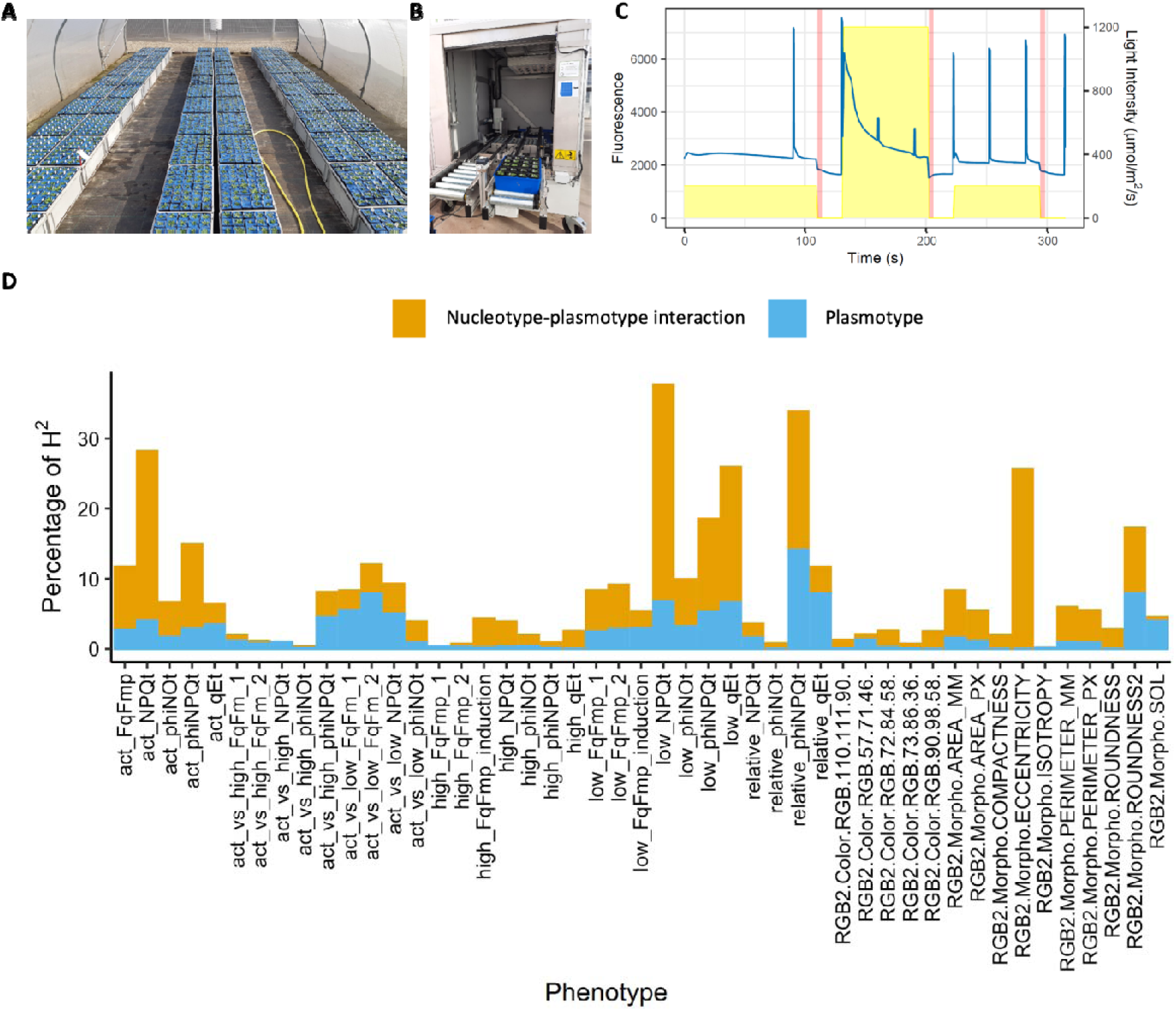
Overview of the experimental design of the cybrids grown in a gauze tunnel, the phenotyping system and H^2^ of the measured phenotypes. A) Image of the gauze tunnel experiment in spring 2022. B) Image of the high-throughput chlorophyll fluorescence phenotyping system. C) The 6-minute light regime and chlorophyll fluorescence measuring regime used to phenotype the cybrid panel. The blue line shows the fluorescence signal of one plant, in response to the light conditions (on the right y-axis). The peaks represent measurements that are used to calculate F’ immediately by Fm’’. The application of far-red light is indicated by the red lines. D) Percentage of H^2^ explained by the plasmotype (blue) and nucleotype-plasmotype interaction (orange) for phenotypes measured over two consecutive years. Ten phenotypes with H^2^ values lower than 5% are not displayed, as fractions of low H^2^ can be inflated. Morphological phenotypes start with “RGB”.

### Plasmotypes causing phenotypic effects

Next, we focused on revealing the plasmotypes that caused phenotypic differences. Using PC analyses, we revealed plasmotypes with deviating effects sizes from the 1,986 phenotypes from the DEPI system including 376 phenotypes from the Phenovator system and 57 phenotypes from the semi-protected experiments. The PCs are calculated for the overall plasmotype effects (i.e., regardless of the nucleotype), and the nucleotype-plasmotype interactions (Figure 4). Based on photosynthetic parameters, the PC show that the relationship between plasmotypes and nucleotypes changes completely between the three different experiments (Figure 4B, 4D and 4F). For example, the Bur-0 plasmotype is separated from the others in the plasmotype PC analysis of the semi-protected experiment (Figure 4E and Supplementary Figure 10), but is grouping centrally in the PCs of the other two experiments (Figure 4A and 4C). The Bur-0 plasmotype was previously found to cause significant phenotypic effects (43), and subsequent in-depth analysis further confirmed a persistent photosynthetic effect under fluctuating light conditions (58). In the fluctuating light conditions of the DEPI experiment, the Bur-0 plasmotype effect can indeed be distinguished (Supplementary Figure 11), although the PC analysis did not identify it as one of the outliers (Figure 4A). This means that environmental conditions strongly influence the differences between PC analyses, emphasizing the strong plasmotype-nucleotype-environment interaction.

**Figure 4.**
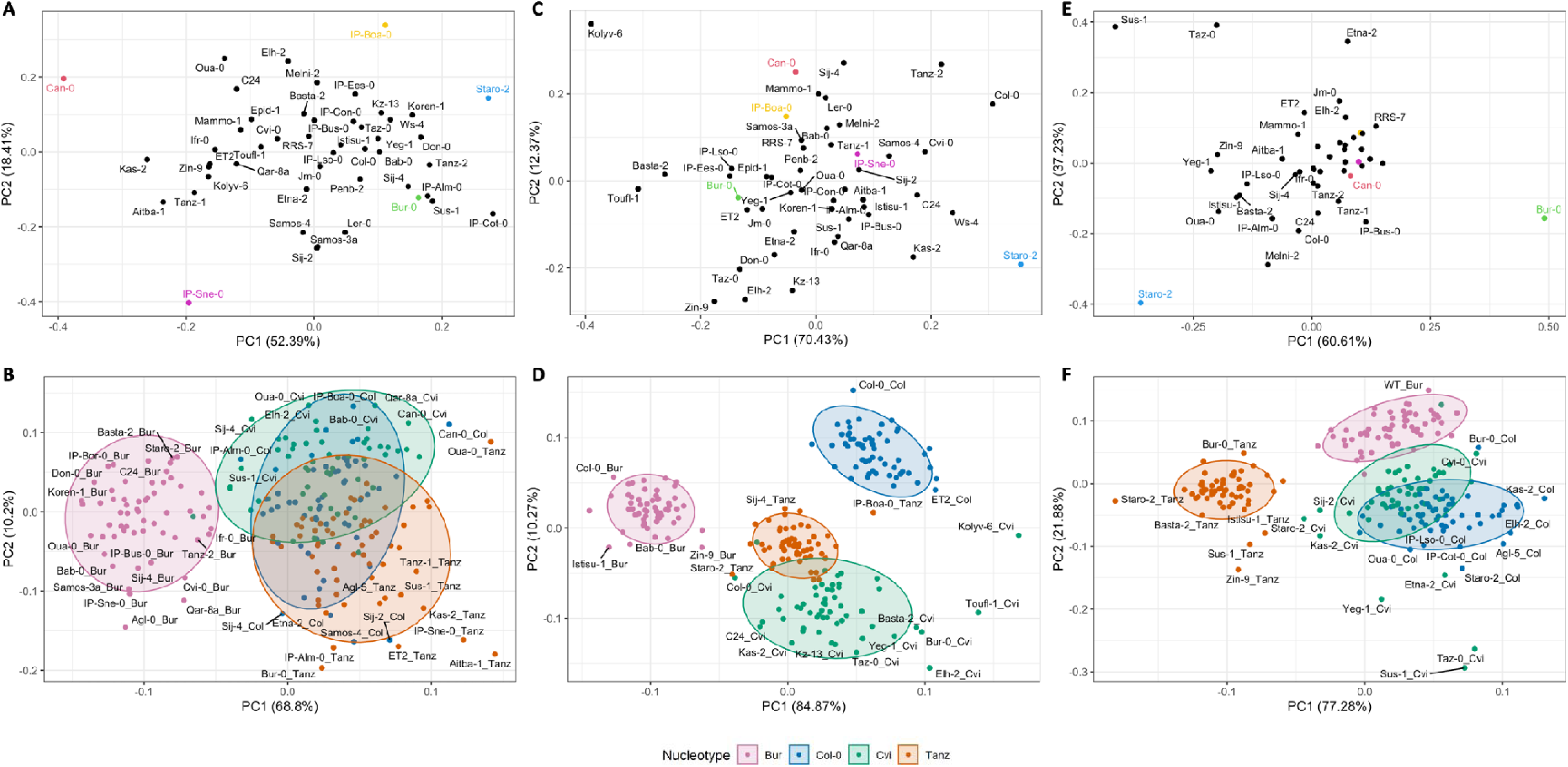
Principal component analysis of the photosynthetic phenotypes in three different experiments. A and B) PC analysis of the photosynthetic phenotypes in the DEPI experiments. C and D) PC analysis of the photosynthetic phenotypes Phenovator experiments. E and F) PC analysis of the semi-protected experiments. Panels A, C and E show the additive analysis, and the outlying plasmotypes are coloured. Panels B, D and F show the analysis of the cybrids separately, and are colour coded per nucleotype. The ovals represent a multivariate t-distribution. Cybrids in panels B, D and F are annotated as “plasmotype_nucleotype”. For all PCs, only the data for photosynthetic parameters are used. In panels A, C and E, the outlying plasmotypes are coloured; these plasmotypes are discussed in the text.

The Bur-0 plasmotype effect showed to be environmentally dependent, however the PC analyses revealed specific plasmotypes to have stronger overall phenotypic effects. The Staro-2 plasmotype effect stands out most persistently in all three experiments (Figures 4A, 4C and 4D). The Staro-2 plasmotype is consistently showing increased Φ_PSII_ in primarily stable low-light conditions (200 µmol m^-2^ s^-1^), as can be seen in the Phenovator experiment (Supplementary Figure 12). Other plasmotypes convey a less consistent phenotypic effect, and are limited to specific environmental conditions. The plasmotypes of IP-Sne and IP-Boa are responsible to the high fraction of H^2^ in *q*_E_ in fluctuating light combined with low temperatures (Figure 2D) when the PC analysis focusses on just these conditions (Supplementary Figure 13). The differences in *q*_E_ are primarily caused by changes in the basal dissipation parameter φ_NO_ (58; Supplementary Figure 13). IP-Sne reduces φ_NO_ by 10.5% in comparison to the average and IP-Boa increases φ_NO_by 11.3% in comparison to the average (Supplementary Figure 14). Both accessions originate from the Iberian Peninsula and they are found in the same subcluster of the neighbour joining tree, but they respond in opposite manners in terms of φ_NO_. This implies that even closely related plasmotypes can cause opposing phenotypes.

The nucleotype-plasmotype interaction explains a higher fraction of H^2^ compared to the additive plasmotype effect. The PC analyses show different plasmotypes as deviant when comparing the four nucleotypes (Figure 4B, 4D and 4F). For example, the Oua-0 plasmotype is only deviating when combined with the Tanz-1 nucleotype, whereas the Agl-0 plasmotype is only deviating when combined with the Bur-0 nucleotype (Figure 4B). Furthermore, while the PC analysis of additive plasmotype effects suggests that Can-0 has large overall phenotypic effects, its position within each nucleotype distribution varies (Figure 4A and 4B). Can-0 is obviously deviating when combined with the Col-0 nucleotype, but less so when combined with of the Bur-0 and Cvi-0 nucleotypes. For the Tanz-1 nucleotype, on the other hand, Can-0 is found to group centrally. This emphasizes the role of nucleotype-plasmotype interactions in determining photosynthetic phenotypes.

### Plasmotype Association Studies

Overall, there is considerable plasmotypic variation conveying photosynthetic differences, but it is generally associated with small effect sizes. This makes the detection of true-positive plasmotype effects complicated. Moreover, methods to narrow down on the causal genetic variations in the plasmotypes are largely lacking. For both cases the concept underlying genome wide association studies (GWAS) can be useful. However, GWAS is developed for mapping in the nucleotype and depends on recombination, which is absent in the plasmotypes. Therefore, we explored an alternative method, which we termed Plasmotype Association Studies (PAS), to avoid confusion with GWAS. In PAS a variant that is unique to a plasmotype or group of plasmotypes with a significant phenotypic effect will be identified. To test this approach, we performed PAS for traits at timepoints where the Bur-0 and Can-0 plasmotypes were found to differ phenotypically. This revealed unique genetic variants for the Can-0 and Bur-0 plasmotypes (Figure 5A and 5B). Theeuwen *et al*. (2022) found a single nucleotide polymorphism (SNP) within the *NdhG* gene of the Bur-0 plasmotype to cause the photosynthetic effect. This causal SNP is amongst the variants with significant associations with the phenotype, thus showing the potential of PAS.

**Figure 5.**
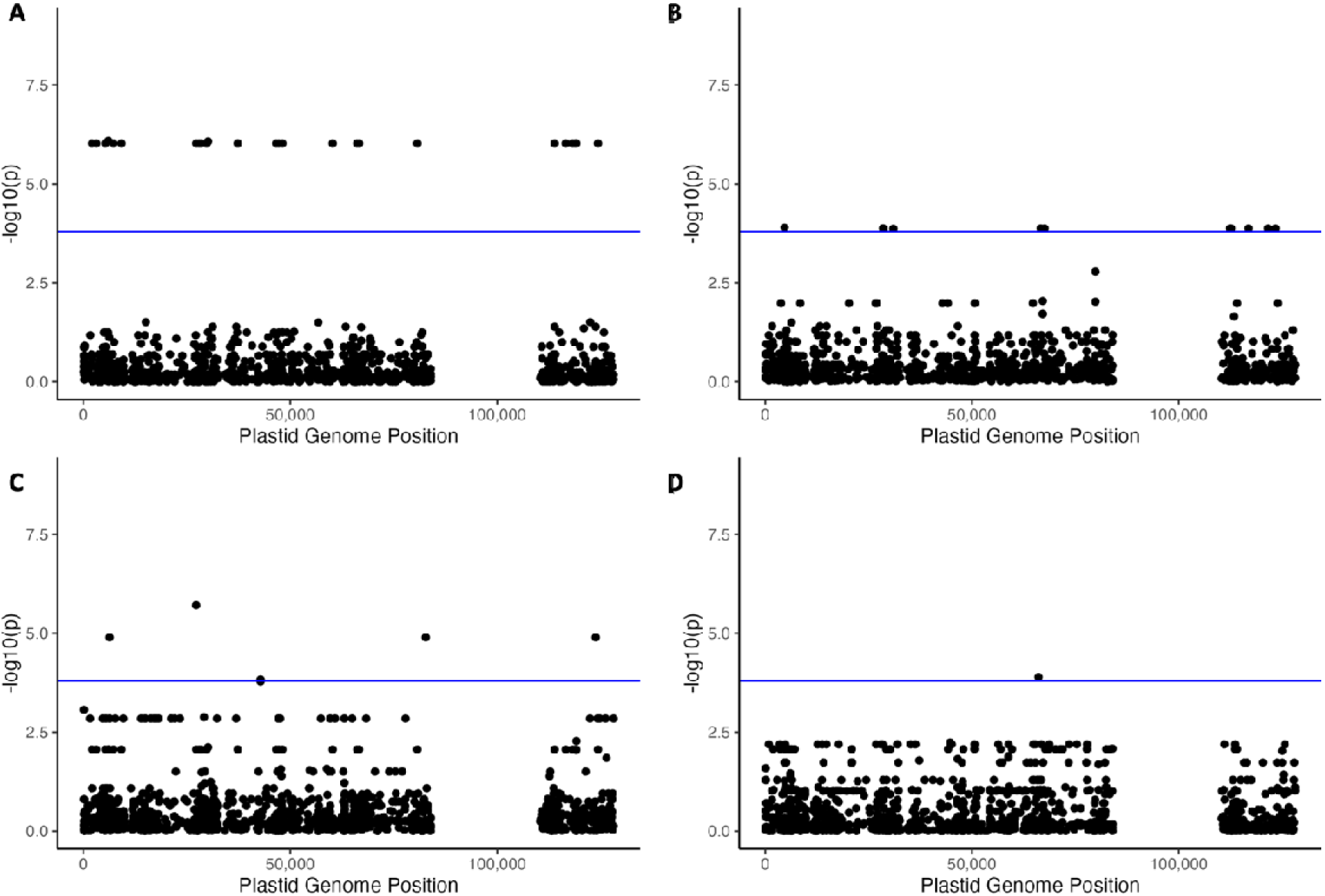
Plasmotype Association Studies (PAS) performed for specific phenotypes acquired in the DEPI experiment. A) LOD-score plot for Φ_PSII_at 55.01 h, revealing variants with LOD-scores exceeding the Bonferroni corrected threshold (blue line), unique to the Bur-0 plasmotype. B) LOD-score plot for Φ_NPQ_ at 78.71 h, revealing a series of significant variants unique to the Can-0 plasmotype. C) LOD-score plot for q_E_ at 127.53h, revealing a significant variant at position 27256 of the chloroplast genome that is shared between IP-Sne and Sij-2. D) LOD-score plot for NPQ at 34.7261h, revealing a significant variant at position 66114 of the chloroplast genome that is shared by Kas-1, Kas-2 and Etna-2. The timepoints mentioned here match the light conditions in Figure 2A. The Bonferroni threshold was determined based on the total number of unique combinations of variants, which is 313 for the chloroplast genomes of the 60 plasmotypes. The resulting Bonferroni threshold (α = 0.05) is therefore set at 3.8.

For phenotypic effects caused by a variant that is shared between several plasmotypes, PAS has increased statistical powers as compared to pairwise comparisons between plasmotypes. Using PAS we found two groups of plasmotypes, each with one shared chloroplast SNP, associated with phenotypic differences. The plasmotypes of IP-Sne and Sij-2 revealed a SNP at base pair position 27,256, a variant that is associated with differences in *q*_E_, NPQ, NPQ_(t)_, *q*_L_, *q*_E(t)_ and NPQ_(t)_ in fluctuating light conditions during low temperatures (Figure 5C). The other group with the plasmotypes of Kas-1, Kas-2 and Etna-2 share a SNP at position 66114, associated with differences in NPQ in high light conditions (Figure 5D). Variants at positions 27,256 and 66,114 are upstream mutations in *rpoC1* and *PsbJ,* respectively. Determining whether these are indeed causal for the phenotypic differences observed requires further experimentation, but it demonstrates the potential power of PAS, as these groups of plasmotypes did not show up when plasmotype effects were examined one by one.

## Discussion

Previous studies using *A. thaliana* have shown there is genetic variation among organellar genomes. However, these studies were based on a limited number of accessions and genetic loci (33, 35, 60, 61). A recent systematic phylogenetic analysis using whole-genome sequencing data, confirmed widespread organellar variation (36). We extended this analysis to include more accessions and used the updated version of the mitochondrial reference genome (21). The updated mitochondrial reference genome is based on Col-0 instead of C24 in the earlier reference genome versions. This ensures that chloroplast, mitochondrial and nuclear diversity can all be compared among accessions. This dataset allowed us to study the characteristics of organellar evolution and diversity at a species level.

We show that the neighbour-joining trees of the mitochondria and chloroplast genomes are highly correlated, and there are no accessions that are assigned to different clusters when comparing the chloroplast and mitochondrial neighbour-joining trees. This means that both organellar genomes have been jointly inheritable in the *A. thaliana* species. Thus paternal transmission of organelles has not affected organellar inheritance in *A. thaliana*, even though paternal transmission of chloroplast may occur at a low frequency (62). Furthermore, we found that the genetic diversity in the chloroplast genomes was eight times that in the mitochondrial genome. This is likely caused by a difference in mutation rate, meaning that the mutation rate difference between the chloroplast and mitochondria is more than twice as high as previously reported (63, 64). The difference between the mutation rate is hypothesized to result from a more efficient repair mechanism in the mitochondria (51). In contrast to the mutation rate being higher in the chloroplast, chloroplast genes appear to be more conserved than the mitochondrial genes. However, this is based on relatively few mutations in the mitochondria due to the low mutation rate. Amongst the 1,531 accessions, we found 1,495 unique plasmotypes and deep branching between subclusters in the neighbour joining tree. Together the mutation accumulation and lack of recombination resulting from the absence of paternal transmission has resulted in diverging plasmotypic subclusters. Therefore, we conclude that there is a high degree of genetic variation between *A. thaliana* plasmotypes, and that most of the genetic diversity is present among the chloroplast genomes.

Organellar genomes are thought to be under purifying selection. While this does not exclude positive selection, organellar variants have long been considered neutral (23). Organellar variation that is adaptive would result in specific plasmotypes confined to specific environmental conditions. An example of this is the spread of the *PsbA* mutation conferring resistance to the herbicide triazine along British railway lines (30). The main plasmotype clusters of the 1,531 *A. thaliana* accessions are spread through most of West Asia and Europe, suggesting that these plasmotypes were not specifically adapted to their environment. This spread is in line with the postglacial recolonization from Southern European refugia, especially from the Balkans (65). Some smaller subclusters are more geographically bound to specific regions, such as a few subclusters from Sub-Saharan Africa. In the Southern Altai mountains in Central Asia a different subcluster is observed than in the Northern Altai mountains. Similarly, several regions within the Atlas Mountains in North Africa are native to only one subcluster. These observations could indicate there is selection for local adaptation to specific environments, though this can also result from genetic drift.

To investigate the impact that organellar genetic variation can have on phenotypes at a species level, we made cybrids with 60 diverse plasmotypes, which is a considerable expansion of the previous cybrid panel we made, containing the reciprocal cybrids of seven accessions (43). This study and another one found that nucleotype–plasmotype interactions determined more variation than plasmotypes alone (43, 66). We therefore now combined the 60 plasmotypes with four diverged nucleotypes, to capture as much nucleotype-plasmotype diversity. Flood *et al.* (2020) found that the plasmotype accounted for 2.9% of the H^2^ and the nucleotype–plasmotype interaction accounted for 5.2% (43). Here, we find on average 1.3% and 2.0% of the variation that can be explained by the plasmotype respectively the nucleotype–plasmotype interaction. Since all seven plasmotypes used in the previous cybrid panel (43) are also included here, the lower percentage of explained H^2^ is either due to the larger genetic variation in the four nucleotypes or due to the wider range of environmental conditions to which the new cybrid panel was exposed. Overall, we find that the plasmotype and nucleotype–plasmotype interaction accounted on average for 3.3% of the total H^2^ of the phenotypes investigated here, while the organellar genomes together make up only 0.4% of the *A. thaliana* genome. Under specific environments, this can go up to 37.9% of the H^2^ to be explained by the plasmotype and 36.6% by nucleotype-plasmotype interaction. As also the PC analysis showed that several plasmotypes and nucleotype-plasmotype interactions contribute to phenotypic differences, we conclude that organellar genetic variation can play a substantial role in a plants environmental adaptation.

In the current study, we primarily show the impact of organellar genetic variation on photosynthesis parameters, and we assume that any differences in photosynthetic phenotypes are most probably associated with chloroplast genes. Nevertheless, also mitochondrial genes are known to play a role in photosynthesis (67). In the new cybrid panel, we found that the Staro-2 plasmotype induced cytoplasmic male sterility (CMS) due to a partially duplicated mitochondrial ORF (68). CMS is often associated with reduced mitochondrial respiration (69, 70). A previous study showed that genotypes displaying CMS also suffer from reduced photosynthetic performance (71). Remarkably, the Staro-2 plasmotype showed the most consistent and highest positive effect on Φ_PSII_of all plasmotypes, suggesting there is a correlation between mitochondrial respiration and photosynthesis. Further analysis of the Staro-2 plasmotype could provide new insights on how reduced mitochondrial respiration can result in increased photosynthetic performance.

Upon closer examination of specific plasmotypes that caused photosynthetic differences, several other insights on the phenotypic impact of plasmotype variation were revealed. The parameter that showed the largest fraction of H^2^ explained by the plasmotype is *q*_E_. As *q*_E_ is a photoprotection mechanism induced under fluctuating light conditions (72), plasmotype variation for *q*_E_ may play a substantial role in light adaptation. One of the plasmotypes influencing the H^2^ for *q*_E_ is the Bur-0 plasmotype. It is found to have faster recovery of Φ_PSII_ than the Col-0 plasmotype after a high-light to low-light transition (58). Even though this plasmotype results in a substantial impact on overall photosynthetic performance and biomass accumulation, it is not one of the consistently deviating plasmotypes in the current study. Can-0 is the plasmotype with the biggest impact on *q*_E_. Different from the Bur-0 plasmotype, the Can-0 plasmotype increases *q*_E_ and decreases Φ_PSII_. The Bur-0 effect is caused by allelic variation of the *NdhG* gene (58), encoding for a subunit of the NDH-like complex. Can-0 has mutations in *NdhF* and *NdhD* (in addition to a variant in *Rps15*), suggesting the importance of the NDH complex in *q*_E_ and Φ_PSII_. The NDH complex is involved in one of the pathways for cyclic electron transport, an essential mechanism in balancing ATP and NADPH production (73). With 12 of the 85 protein-coding genes in the chloroplast encoding components of the NDH complex, variation in these genes may explain a substantial part of the observed photosynthetic differences.

Revealing the genetic variation underlying phenotyping differences remains difficult due to the small effect sizes and multiple testing corrections needed. Statistical power can be increased by using variants that are shared by all plasmotypes showing a phenotypic effect. For this we propose to use a GWAS approach on the plasmotypes, here termed PAS. PAS is similar to GWAS, but as organellar genomes do not recombine like the nuclear genomes, the resulting associations are not subject to linkage disequilibrium decay (74). Consequently, upon PAS it is not possible to identify which of the variants with the same LOD score could be the cause of any observed phenotypic effect. This approach resembles how GWAS has been used to associate genetic variants with phenotypic differences in bacterial genomes (75–77). We performed PAS only on the chloroplast genome due to the dynamic properties of the mitochondrial genome and heterozygosity of some variants, making correct associations difficult (53, 54). Nevertheless, significant associations found in the chloroplast may be caused by genetic variation in the mitochondrial genome, due both organellar genomes of a plasmotype inheriting together. Minor allele frequency thresholds are used in GWAS to avoid the chance of identifying false-positive associations between alleles and phenotypic variation. Due to the small population size of our cybrid panel and the high degree of genetic diversity a minor allele frequency threshold of 5% excludes most variants. Nevertheless, several significant associations are identified. These associations will need to be experimentally confirmed to identify the causal variant. For organellar genomes, transformation approaches would be the most promising (78–82). Although, at the moment, most of the methods that allow changing any genetic variant or generating knockouts in all copies of the organellar genomes are still under development. An alternative would be to use a genetic exclusion approach, where plasmotypes with shared variants are phenotyped for the same trait. In the absence of a common phenotypic difference among these variants compared to the rest, that specific variant can be excluded (58). Overall, we show that PAS has the potential to identify candidate alleles associated with phenotypic effects.

Previous research already showed that different phenotypes can arise due to plasmotypic variation (23, 43, 66, 83), but a systematic analysis on how plasmotypic variation contributes to photosynthetic performance was lacking. Here, we used *A. thaliana* as a model species to conclude that plasmotype diversity is widespread and that several plasmotypes harbour variants that can significantly impact photosynthetic performance. The contribution of the plasmotype variation as part of total phenotypic variation is relatively small, however it can be substantial especially in dynamic environmental conditions. Currently, the range of crops in which cybrids can be made via haploid induction is quickly expanding (84–87). Given the widespread plasmotypic effects on photosynthetic performance and the advancing capabilities to produce cybrids, the time has now come to tap into organellar diversity for crop improvement.

## Material and Methods

### Analysis of *A. thaliana* organellar diversity

The recalling of the variant was done using publicly available sequencing data. The data were obtained from ENA for the following datasets; 1135 worldwide accessions (PRJNA273563) (45), 117 Chinese accessions (PRJNA293798) (48), 75 African accessions (PRJEB19780) (46), 14 Madeiran accessions (PRJEB23751) (49), 1 Tibetan accession (47), 7 global accession (PRJEB29654) (43), 192 Dutch accessions (88) and 4 related species (*A. lyrata*, *A. halleri*, *A. carpatica* and *Capsella rubella*). The variant calling was done with the same pipeline as described in (43). One accession was removed due to a missing .fastq file. All accessions were mapped to the *A. thaliana* Col-0 reference genome (TAIR10.1, GCA_000001735.2)(21). To remove probable false-positive calls, the organelle genome variants were filtered based on their quality-by-depth score (QD). The chloroplast call set was filtered with a QD of minimally 25, leaving 4095 variants and for the mitochondrial call set was filtered with a QD of minimally 20, leaving 1152 variants. Ten accessions with more than 50% of the chloroplast genome variants called heterozygous were removed.

Subsequent analyses were done in R (version 4.0). The four related species were left out of the analysis, unless stated explicitly. Ape (version 5.4-1) (89) and vcfR (version 1.12.0) (90) were used to perform hierarchical clustering, based on complete-linkage clustering. The dendogram was cut using a k=20 as an arbitrary cut-off, but based on the elbow method. Principal component analyses where done with SeqArray (version 1.28.1) (91) and SNPRelate (version 1.22.0), as these packages allowed to include multiallelic variants (92). Geographic locations were taken from the respective papers, and plotted on the map using rworldmap (version 1.3-6) and rworldxtra (version 1.01). Figures where made using ggplot2 (version 3.3.2) and GGally (version 2.0.0).

### Plant material and creation of novel cybrid panel

The wildtype plants were either present in the laboratory, obtained via the Nottingham Arabidopsis Stock Centre or send to us via colleagues (Supplementary Table 3). We used the *GFP-tailswap* line as described in Flood *et al.* (2020), as haploid inducer line. The cybrid panel was made as described in Flood *et al.* (2020). For this study that meant the *GFP-tailswap* line was crossed as paternal line to all 60 accessions, and the F_2_ genotypes homozygous for *GFP-tailswap* were selected. This resulted in 60 *GFP-tailswap* genotypes, all having a different plasmotype. Subsequently, these 60 *GFP-tailswap* genotypes were crossed to the four nucleotype donors, with the *GFP-tailswap* as the maternal parent. 14 cybrids in the new panel were overlapping with a previous cybrid panel, and were obtained from Flood *et al.* (2020). All cybrids were propagated in the same controlled greenhouse, to exclude possible batch effects.

### Genotyping of novel cybrid panel

DNA extraction was done in 96 deep well plates. Single rosette leaves were harvested from individual plants and placed in the deep well plates, snap frozen in liquid nitrogen and ground with a Retsch MM300 TissueLyser. 100 mL extraction buffer (200 mM Tris-HCl, 25 mM EDTA and 1% SDS) was prepared by adding 40 µL of 20 mg/mL RNase A. 500 µL of extraction buffer including RNase A was added to each well, and incubated at 37 °C for 1 hour, and inverted every 15 minutes. To pellet the debris the plates were centrifuged for 5 minutes at 3000 x g. In a new deep well plate 130 µL KAc buffer (98.14 g potassium acetate and 3.5 mL Tween were added to 160 mL H_2_O and H_2_O was added to reach 200 mL) and 400 µL lysate were added. The plates were sealed and inverted for 2 minutes, and incubated on ice for 10 minutes. To pellet the debris the plates were centrifuged for 5 minutes at 3000 x g, and 400 µL of supernatant was transferred to a new plate containing 440 µL Sera-Mag Speedbeads (Cytiva Europe) diluted in PEG buffer. Plates were sealed and place on a shaking table for 30 minutes. Next, the plates were placed for 5 minutes on a magnet and the supernatant was removed and washed with 500 µL 80% EtOH, and repeated three times. The plates were left to evaporate in the fume hood for 1 hour and the DNA was resuspended in 50 µL milliQ.

Consecutively the Hackflex protocol was used to do the library preparation (93). Samples were pooled, and the fraction showing strands of roughly 300 to 500 bp in size were selected and sequenced for on average 8X whole genome coverage sequencing by Novogene (UK) Ltd. The reads were trimmed using Cutadapt (94), removing the adaptor sequences and bases from the 5’ and 3’ ends with a Phred quality score below 20. Reads shorter than 75 bp were also removed. The reads were then mapped to the *A. thaliana* Col-0 reference genome, TAIR10.1, using speedseq and the Burrow-Wheeler aligner, BWA-MEM (95, 96). The alignments were sorted and indexed using Samtools (97). Duplicate reads were marked using GATK (98). Variant calling was performed using freebayes and the resulting variant call format (VCF) file was separated into three VCF files – one for each of the nuclear, mitochondrial and plastid genomes (99). Distance matrices were calculated for each of the three genomes using PLINK (100), and the ape package in R was used to produce neighbour-joining trees (101). The cybrids with incorrect genotypes were removed from all statistical analyses.

### Phenotyping and data analysis

#### Semi-protected conditions experiments

Plant growth took place in an outdoor, gauze covered tunnel at Unifarm, Wageningen University and Research, the Netherlands (51.9882583, 5.66119897). The base of the tunnel measured 8 × 5 m and was enclosed by synthetic gauze material that was largely penetrable by rainwater, sunlight and wind so to provide conditions similar to those encountered in the field. Rain gauges placed inside and outside the tunnel confirmed that all rainwater was able to penetrate the gauze. Light irradiance was measured both inside the tunnel and at a metrological station within 500 m from the tunnel (Supplementary Figures 15 and 16). A comparison of readings from both locations, during the growing period of spring 2020, indicated that the gauze decreased the light irradiance that penetrated through to the growing area on average by 94.3 µmol m^-2^ s^-1^. The floor of the tunnel was covered in black landscape material and the tunnel contained a zipper door to prevent the entry of any undesired interferents.

Black plastic pots measuring 7×7×18 cm (Bestebreurtje B.V., Huissen, the Netherlands) were used for individual plants as they allowed sufficient depth for unlimited root development. Grey plastic trays measuring 40×60×20 cm were used to hold 40 of the aforementioned pots. The trays were organized in five rows of 11 trays and one row of 13 trays. A small seedling tray was placed in the bottom of each large grey tray to raise the pots above the edge of the grey tray. The plant pots were filled with a mixture of 40 % sand and 60 % peat provided by Lensli Substrates (Katwijk, the Netherlands). The substrate includes YARA PG MIX™ which contains 15-10-20+3 of N, P_2_O_5_, K_2_O and MgO. The added fertilizer is in powder form and results in complete substrate values of 1.0 and 5.7 for electrical conductivity and pH, respectively. No additional nutrition was applied during the experiment.

In spring of 2020 plastic wrapping was placed over the trays during the night for the first 14 days of growth to protect from cold temperatures. The same was done in spring 2021 for the first five days of growth. In spring 2020 grey rubber covers were placed on each pot after seedlings had established, leaving 0.5 cm space for water to reach the soil. In spring 2021 the soil was covered with blue Friedola Mega Stop mats as described in (102). The pots were evenly watered as needed according to weather conditions and rainfall. Anti-slug/snail pellets were placed in small piles on the ground around the perimeter of the tunnel. Remote sensors from 30MHz (Amsterdam, the Netherlands) monitored and recorded light irradiance and temperature every minute at plant level.

Phenotyping was primarily done using the high-throughput phenotyping system PlantScreen^TM^ SC System provided by Photon Systems Instruments spol. s r.o (Drásov, Czech Republic). Using the chlorophyll fluorescence camera and an RGB camera, we obtained a range of photosynthetic and morphological parameters. A custom made 6-minute fluctuating light protocol (Figure 3C), with 5 F’ and Fm’ measurements, allowed us to calculate Φ_PSII,_ Φ_NOt,_ Φ_NPQt_, NPQt and *q*_E_ at different moments during the 6-minute protocol. A custom R script converted these measurements into 37 parameters capturing the dynamic response to the fluctuating light protocol. A custom mask using the Schedular software was generated to phenotype 20 plants, with a 7 x 7 cm grid size. In spring and autumn 2020, the grey rubber plates were complemented with additional rubber strips, to mask soil-grown algae being registered. In spring 2021 the blue mats ensured the algal growth was not recorded. Automatic masking by the Data Analyser software ensured background noise was removed, and the plant mask was generated. The RGB camera generated an additional 20 parameters quantifying the morphological characteristics and colour properties. In spring 2020 the shoots were also harvested to measure the shoot dry weight when the first flower of a plant opened.

During spring 2020 221 cybrids were ready and could be sown. The cybrids were sown in an unbalanced, incomplete block design to randomize the cybrids among the pots meaning the nucleotypes being randomized among the trays and the plasmotypes being randomized within the trays. The number of replicates ranged between 10-12 for the cybrid genotypes and 60-80 for the four wildtypes. Due to the number of pots, sowing was spread over three days, with cybrids with the Bur-0 nucleotype being sown on March 18^th^ 2020, cybrids with Col-0 and Cvi nucleotypes on March 19^th^ and cybrids with Tanz nucleotypes on March 20^th^. Approximately 4 pre-germinated seedlings were sown per pot, and all but the healthiest seedling was removed 20 days after sowing. The trays containing nucleotypes Cvi-0, Col-0, Bur-0 and Tanz-1 were phenotyped in the PlantScreen^TM^ system at 35, 38, 38 and 38 days of growth, respectively.

During spring 2021 all 240 cybrids were ready and could be sown. A complete randomized block design was used, in a way that all 240 cybrids were sown once every six trays. Such six trays were arranged in a 2 x 3 orientation, and together formed one big block, with the total experiment having n=12. As phenotyping in the PlantScreen^TM^ system allowed only 20 plants, we ensured that all cybrids within a block were measured back-to-back to correct over the day and time as best as possible. Again, the sowing was split over three days, cybrids with the Bur-0 nucleotype were sown on March 17^th^ 2021, cybrids with the Col-0 nucleotype on March 18^th^, and cybrids with the Cvi and Tanz nucleotype on March 19^th^. After 24 days of growth, all but the healthiest seedling were removed. Phenotyping in the PlantScreen^TM^ system happened on May 3^rd^ and 4^th^, meaning the plants ranged between 45 and 48 days old.

Growth measurements, red-green-blue (RGB) measurements and chlorophyll fluorescence parameters (Φ_PSII_, Φ_NPQ_, Φ_NO_, NPQ and *q*_E_) from the two seasons were compiled. Plants that did not establish well were removed via outlier detection on the basis of leaf area. This was done by calculating the mean leaf area per genotype and individuals with a leaf area less than the mean – 1.5 × standard deviation were removed. Plants with the Ely plasmotype were removed from all subsequent analyses as this plasmotype confers a particularly large reduction in Φ_PSII_, thus obscuring smaller contributions of the plasmotype to the heritability of photosynthetic traits (43). Linear mixed models were constructed for each response variable using the lme4 package in R (version 4.1.0) (103). The contribution of the nucleotype, plasmotype and the nucleotype-plasmotype interaction to each phenotypic variance was quantified by the variance components for these model terms in the random model (1), using a restricted maximum likelihood approach.

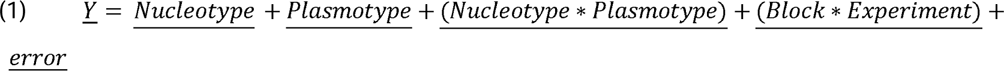

The broad sense heritability (H^2^) was calculated as the sum of the genetic variance components (Nucleotype, Plasmotype and Nucleotype × Plasmotype) relative to the total phenotypic variance. The contribution of each genetic component to the H^2^ was summarized as a percentage of the total H^2^. Phenotypes with a H^2^ lower than 5% were removed from all subsequent analyses, as variation in these phenotypes cannot be accurately predicted based on genotype (43).

To investigate the main effect of plasmotype and its interaction with nucleotype, a mixed model was fitted with fixed main effects for nucleotype and plasmotype and their fixed interaction and random terms for blocks and error. This resulted in the model given in equation (2).

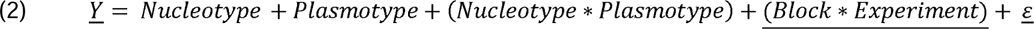

#### DEPI experiment

Phenotyping screens were carried out using the Dynamic Environmental Photosynthetic Imaging (DEPI) system (56), with modifications as described in (104). The DEPI system allows all of the plants to be measured simultaneously, allowing repeated measurements under fluctuating light conditions. The plants were grown in climate controlled conditions for 18 days and moved into the DEPI system to acclimate for three days. The DEPI experiment started (t = 0 h) at midnight on day 21. Two different light regimes were used, one day the light intensity was stable at 200 µmol m^-2^ s^-1^ and the other day it was fluctuating light, with sinusoidial increases and decreases over the day. These fluctuations were characterized as 18 minutes low light, 2 minutes darkness, 8 minutes high light and 2 minutes darkness (see for the full description Cruz *et al.*, 2016). In total we ran five DEPI experiments, each with 224 plants. Using an incomplete block design, the cybrids were distributed so that they were sown in four out five experiments (n = 4). Every experiment had 8 replicates of the four wildtype parents.

Growth measurements and chlorophyll fluorescence parameters (F_v_/F_m_, NPQ, NPQ_(t)_, Φ_PSII_, Φ_NPQ_, Φ_NO_, *q*_E_, *q*_E(t)_, *q*_I_, *q*_I(t)_, *q*_L_) from five experiments were compiled. To remove plants that performed poorly in comparison to the other individuals of the same genotype outlier detection was done on the dark-adapted yield of PSII photochemistry (F_v_/F_m_). The mean F_v_/F_m_ for all plants was calculated and plants with an F_v_/F_m_ less than the mean – 1.5 × standard deviation were removed. As in the semi-controlled experiments, plants with the Ely plasmotype were removed from all subsequent analyses.

Statistical modelling was carried out as with the data from the semi-controlled experiments, using equation (3) for the estimation of variance components and the calculation of H^2^ and equation (4) for the calculation of the main effects. Phenotypes with a H^2^ lower than 5% were removed for all subsequent analyses.

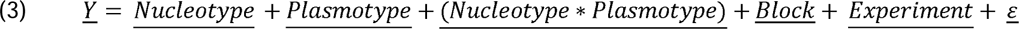

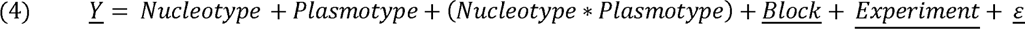

#### Phenovator experiment

The cybrid panel was phenotyped using the Phenovator, a high-throughput phenotyping platform in which 1440 *A. thaliana* plants can be grown (57). Plants have been measured seven times a day for photosystem II efficiency and growth, except during the three days with fluctuating light. On fluctuating-light days the plants have only been measured once, on the middle of the day. The plants were grown with a 12 hour photoperiod (Supplementary Figure 8). For the first 17 days light intensity was set at 200 µmol m^-2^ s^-1^. On day 18 fluctuating light conditions were started for three days. During these fluctuating light regimes, every twenty minutes the light switched between 500 µmol m^-2^ s^-1^ and 100 µmol m^-2^ s^-1^, except during measurements. Only one measurement was performed during these days, during one hour. At the end of day 21, the temperature was decreased to from 20 °C to 12 °C during the day, and from 18 °C to 6 °C in the night. Throughout the experiment the relative humidity was kept at 70%. Plants were grown on a 4x4x4 cm rockwool substrate provided by Grodan B.V. (Roermond, the Netherlands), and irrigated weekly with a nutrient solution (Supplementary Table 3). The cybrid panel was screened using this system four times. One run included only the cybrids with the Col-0 nucleotype, and a randomized complete block design was used (n=24). The other three runs included the full cybrids panel, and was sown in a randomized complete block design (n=6 in each run).

Leaf area and Φ_PSII_ measurements were extracted from image files produced by the Phenovator using custom software (57). Measurements from the four separate experiments were subsequently compiled. Outlier detection was used to remove plants that did not establish well, by removing individuals with a leaf area (15 days after sowing) that was 1.5 × standard deviation lower than the mean per genotype. Furthermore, complete genotypes that showed poor germination, resulting in stunted growth, were removed. This because these cybrids were not always removed by the outlier detection as the whole genotype was stunted. This included all cybrids with the Cvi nucleotype in one experiment and all of the Cvi^Sha^ cybrids. Plants with the Ely plasmotype were removed from all subsequent analyses, as in the semi-controlled and DEPI experiments. Statistical modelling was carried out as with the data from the semi-controlled and DEPI experiments, using equation (5) for the estimation of variance components and the calculation of H^2^ and equation (6) for the calculation of the main effects. Phenotypes with a H^2^ lower than 5% were removed from all subsequent analyses. The model includes image position, to correct for intensity of saturating light as received by each of the twelve positions. Experiment represents the four individual runs.

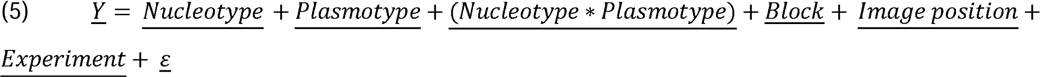

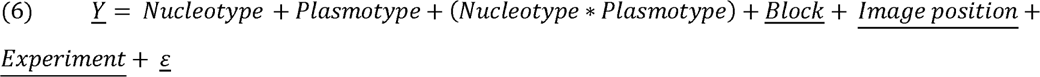

For all the three experiments described above a separate principal component (PC) analysis was carried out with R, using the main effects for each plasmotype. Further PCs were performed for each of the four nucleotypes using the main effects for the nucleotype-plasmotype interactions, which are equal to the Best Linear Unbiased Estimate. These PC analyses were done for the photosynthetic parameters only.

### Plasmotype Association Studies

A Genome-Wide Association Studies (GWAS) method was used to test the association between genetic variants in the plastid genome and the phenotypes measured in the DEPI experiments. Due to the lack of recombination within the plastid genome, the interpretation of the GWAS results deviates from a standard GWAS performed on the nuclear genome. Unless genetic variants are shared between different plasmotypes, all of the SNPs within a plasmotype will be equally associated with a phenotype. Thus, to avoid confusion with a nuclear GWAS, this test has been named a Plasmotype Association Study (PAS). To carry out the PAS, the main effect for each plasmotype from the DEPI phenotyping experiments were used alongside the VCF file for the plastid genome on the basis of 40X sequencing coverage. Binary PED files were produced from the plastid VCF file using PLINK (100). The main effect for each of the 1986 phenotypes were added manually to the .fam file. GEMMA was used to produce a centred relationship matrix for the plastid genomes of the 60 progenitor lines and to run a univariate association study (105).

## Data, code and genetic material availability

The raw data files with genotyping and phenotyping data are available via Zenodo (https://doi.org/10.5281/zenodo.11259864). R scripts and Linux commands for data analyses are available via GitLab (https://git.wur.nl/tom.theeuwen/novel-cybrids.git). Seeds of the cybrids are available upon request.

## Acknowledgements and funding

We thank Andreas Hemp for providing seeds of accession ET2, John Doonan for providing seeds of accession Penbont-1 and Carlos Alonso-Blanco for seeds of Agl-5. We also thank Ben Auxier, Roel van Bezouw, René Boesten, Ramon Botet, Pádraic Flood, Willem Kruijer and Antoine Languilaume for insightful discussions and José van de Belt and Jordy Litjens for their help in the construction of the new cybrids. Ben Auxier is acknowledged for feedback on the manuscript. This work was, in part, supported by the Netherlands Organization for Scientific Research (NWO) through ALWGS.2016.012 (TPJMT).

## Supplementary Tables

**Supplementary Table 1.** H^2^ and fractions of H^2^ explained by the plasmotype and nucleotype-plasmotype interaction for the DEPI experiment. The values are grouped per trait, both over all timepoints as well as per environmental conditions as exposed to.

| Phenotype | Environmental conditions | $H^2$ | Fraction NxP | Fraction P | Fraction N |
| --- | --- | --- | --- | --- | --- |
| fvfm | Stable light day 1 | 3.8% | NA | NA | NA |
| fvfm | Fluctuating light day 1 | 3.2% | NA | NA | NA |
| fvfm | Stable day 2 | 9.3% | 2.9% | 0.0% | 97.1% |
| fvfm | Fluctuating light day 2 | 7.4% | 0.0% | 0.0% | 100.0% |
| fvfm | Stable light cold day | 68.6% | 0.0% | 0.0% | 100.0% |
| fvfm | Fluctuating light cold day | 67.1% | 0.0% | 0.0% | 100.0% |
| growth | Stable light day 1 | 61.4% | 5.1% | 0.5% | 94.3% |
| growth | Fluctuating light day 1 | 61.6% | 5.1% | 0.3% | 94.6% |
| growth | Stable day 2 | 59.8% | 5.8% | 0.1% | 94.1% |
| growth | Fluctuating light day 2 | 61.8% | 6.0% | 0.2% | 93.8% |
| growth | Stable light cold day | 59.7% | 7.0% | 0.0% | 93.0% |
| growth | Fluctuating light cold day | 60.2% | 7.0% | 0.0% | 92.9% |
| npq | Stable light day 1 | 9.2% | 6.7% | 3.7% | 89.6% |
| npq | Fluctuating light day 1 | 12.9% | 1.1% | 1.3% | 97.6% |
| npq | Stable day 2 | 10.6% | 3.6% | 0.0% | 96.4% |
| npq | Fluctuating light day 2 | 15.0% | 1.3% | 0.5% | 98.1% |
| npq | Stable light cold day | 34.6% | 3.6% | 1.7% | 94.7% |
| npq | Fluctuating light cold day | 27.5% | 3.3% | 1.0% | 95.7% |
| npqt | Stable light day 1 | 27.2% | 0.9% | 0.6% | 98.4% |
| npqt | Fluctuating light day 1 | 34.8% | 0.7% | 1.7% | 97.6% |
| npqt | Stable day 2 | 24.9% | 0.0% | 1.0% | 99.0% |
| npqt | Fluctuating light day 2 | 45.9% | 0.1% | 1.4% | 98.4% |
| npqt | Stable light cold day | 30.2% | 0.6% | 0.7% | 98.7% |
| npqt | Fluctuating light cold day | 20.3% | 0.5% | 0.5% | 99.0% |
| phi2 | Stable light day 1 | 15.8% | 1.6% | 0.4% | 97.9% |
| phi2 | Fluctuating light day 1 | 15.4% | 0.2% | 0.8% | 99.0% |
| phi2 | Stable day 2 | 11.9% | 0.8% | 1.7% | 97.6% |
| phi2 | Fluctuating light day 2 | 20.7% | 0.2% | 1.0% | 98.8% |
| phi2 | Stable light cold day | 41.4% | 0.0% | 0.1% | 99.9% |
| phi2 | Fluctuating light cold day | 33.7% | 0.0% | 0.4% | 99.6% |
| phino | Stable light day 1 | 46.1% | 1.6% | 0.8% | 97.6% |
| phino | Fluctuating light day 1 | 55.7% | 2.2% | 1.1% | 96.7% |
| phino | Stable day 2 | 50.0% | 0.2% | 0.2% | 99.6% |
| phino | Fluctuating light day 2 | 64.8% | 1.0% | 0.8% | 98.2% |
| phino | Stable light cold day | 6.9% | 2.8% | 2.0% | 95.2% |
| phino | Fluctuating light cold day | 12.0% | 1.0% | 1.3% | 97.7% |
| phinpq | Stable light day 1 | 19.7% | 1.1% | 0.3% | 98.7% |
| phinpq | Fluctuating light day 1 | 23.0% | 0.2% | 0.9% | 98.9% |
| phinpq | Stable day 2 | 16.8% | 0.2% | 1.5% | 98.3% |
| phinpq | Fluctuating light day 2 | 31.4% | 0.1% | 1.1% | 98.9% |
| phinpq | Stable light cold day | 51.5% | 0.0% | 0.2% | 99.8% |
| phinpq | Fluctuating light cold day | 36.9% | 0.1% | 0.5% | 99.4% |
| qesv | Stable light day 1 | 8.6% | 10.6% | 6.5% | 82.9% |
| qesv | Fluctuating light day 1 | 27.2% | 1.4% | 4.8% | 93.8% |
| qesv | Stable day 2 | 12.7% | 0.0% | 3.4% | 96.6% |
| qesv | Fluctuating light day 2 | 42.2% | 1.3% | 3.5% | 95.2% |
| qesv | Stable light cold day | 15.4% | 8.0% | 2.7% | 89.3% |
| qesv | Fluctuating light cold day | 27.1% | 2.2% | 6.4% | 91.4% |
| qet | Stable light day 1 | 11.0% | 7.2% | 2.6% | 90.2% |
| qet | Fluctuating light day 1 | 30.2% | 1.9% | 3.0% | 95.1% |
| qet | Stable day 2 | 12.3% | 0.6% | 1.9% | 97.5% |
| qet | Fluctuating light day 2 | 48.5% | 1.5% | 1.4% | 97.1% |
| qet | Stable light cold day | 9.0% | 6.8% | 1.7% | 91.5% |
| qet | Fluctuating light cold day | 13.1% | 0.5% | 1.4% | 98.1% |
| qi | Stable light day 1 | 12.3% | 6.2% | 3.4% | 90.4% |
| qi | Fluctuating light day 1 | 15.1% | 0.6% | 0.0% | 99.4% |
| qi | Stable day 2 | 21.0% | 6.2% | 0.0% | 93.8% |
| qi | Fluctuating light day 2 | 14.7% | 2.7% | 0.1% | 97.3% |
| qi | Stable light cold day | 44.3% | 1.6% | 0.4% | 97.9% |
| qi | Fluctuating light cold day | 34.1% | 2.3% | 0.0% | 97.7% |
| qit | Stable light day 1 | 21.8% | 0.8% | 1.0% | 98.1% |
| qit | Fluctuating light day 1 | 30.7% | 0.5% | 1.0% | 98.5% |
| qit | Stable day 2 | 23.4% | 0.5% | 0.8% | 98.8% |
| qit | Fluctuating light day 2 | 26.5% | 0.3% | 1.9% | 97.8% |
| qit | Stable light cold day | 34.3% | 0.5% | 1.5% | 98.0% |
| qit | Fluctuating light cold day | 30.8% | 0.7% | 0.6% | 98.7% |
| ql | Stable light day 1 | 43.3% | 2.4% | 1.4% | 96.2% |
| ql | Fluctuating light day 1 | 46.0% | 3.6% | 1.0% | 95.4% |
| ql | Stable day 2 | 47.4% | 0.3% | 0.2% | 99.5% |
| ql | Fluctuating light day 2 | 54.4% | 1.6% | 0.9% | 97.5% |
| ql | Stable light cold day | 2.3% | NA | NA | NA |
| ql | Fluctuating light cold day | 5.7% | 0.0% | 1.5% | 98.5% |

**Supplementary Table 2.** Nutrient solution as used for growing A. thaliana on rockwool substrate.

|  | Concentration | Unit |
| --- | --- | --- |
| NH<sub>4</sub> | 1.7 | mmol/L |
| K | 4.13 | mmol/L |
| Ca | 1.97 | mmol/L |
| Mg | 1.24 | mmol/L |
| NO<sub>3</sub> | 4.14 | mmol/L |
| SO<sub>4</sub> | 3.14 | mmol/L |
| P | 1.29 | mmol/L |
| Fe* | 21 | μmol/L |
| Mn | 3.4 | μmol/L |
| Zn | 4.7 | μmol/L |
| B | 14 | μmol/L |
| Cu | 6.9 | μmol/L |
| Mo | 0.5 | μmol/L |
| EC | 1.4 | mS/cm |
| pH** | 6.1 |  |

\*Consists of 50% Fe-DTPA 3% and 50% Fe- EDDHSA 3%
\*\*pH adjusted with Potassium hydroxide and Sulphuric acid

**Supplementary Table 3.** Description of 60 accessions used to produce the cybrid panel.

| Accession name | Name in VCF | Stock number | Obtained from | Latitude | Longitude |
| --- | --- | --- | --- | --- | --- |
| Agl-0 | Agl-0 | CS799920 | NASC | 32.97243 | -5.44856 |
| Agl-5 | Agl-5 | - | Carlos Alonso-Blanco | 32.97243 | -5.44856 |
| Aitba-1 | Aitba-1 | CS76649 | NASC | 31.48 | -7.45 |
| Bab-0 | Bab-0 | CS799910 | NASC | 35.04256 | -5.02369 |
| Basta-2 | Basta-2 | CS76692 | NASC | 51.82 | 79.48 |
| Bur | Bur_Cybrid | CS2110195 | Laboratory of Genetics (WUR) | 52.9 | -9 |
| C24 | C24_Cybrid | CS2110196 | Laboratory of Genetics (WUR) | 40.2077 | -8.42639 |
| Can-0 | Can-0 | CS2110034 | Laboratory of Genetics (WUR) | 29.2144 | -13.4811 |
| Col | Col_Cybrid | CS2110197 | Laboratory of Genetics (WUR) | - | - |
| Cvi-0 | Cvi-0 | CS902 | Laboratory of Genetics (WUR) | 15.1111 | -23.6167 |
| Don-0 | Don-0 | CS2110040 | NASC | 36.83 | -6.36 |
| Elh-2 | Elh-2 | CS799925 | NASC | 31.47197 | -7.40644 |
| Elk-1 | Elk-1 | CS799922 | NASC | 32.53516 | -6.014969 |
| Ely | Ely_Cybrid | CS2110198 | Laboratory of Genetics (WUR) | 50.5 | -4.5 |
| Epidauros-1 | Epidauros-1 | CS76844 | NASC | 37.6 | 23.08 |
| ET2 | - | - | Andreas Hemp | 13.153 | 38.4 |
| Etna-2 | Etna-2 | CS76487 | NASC | 37.69 | 14.98 |
| Ifr-0 | IFr-0 | CS799918 | NASC | 33.55006 | -5.17465 |
| IP-Alm-0 | IP-Alm-0 | CS799191 | NASC | 39.88 | -0.36 |
| IP-Boa-0 | IP-Boa-0 | CS799204 | NASC | 40.4 | -3.88 |
| IP-Bor-0 | IP-Bor-0 | CS76717 | NASC | 42.49 | -6.71 |
| IP-Bus-0 | IP-Bus-0 | CS799207 | NASC | 36.97 | -3.28 |
| IP-Con-0 | IP-Con-0 | CS76780 | NASC | 37.94 | -5.6 |
| IP-Cot-0 | IP-Cot-0 | CS799229 | NASC | 41.83 | -5.38 |
| IP-Ees-0 | IP-Ees-0 | CS799235 | NASC | 40.59 | -4.15 |
| IP-Lso-0 | IP-Lso-0 | CS799265 | NASC | 38.86 | -3.16 |
| IP-Per-0 | IP-Per-0 | CS77169 | NASC | 37.6 | -1.12 |
| IP-Piq-0 | IP-Piq-0 | CS799297 | NASC | 42.1 | -2.56 |
| IP-Sne-0 | IP-Sne-0 | CS77258 | NASC | 37.09 | -3.38 |
| Istisu-1 | Istisu-1 | CS75433 | NASC | 38.9786 | 48.5594 |
| Jm-0 | Jm-0 | CS6748 | NASC | 49 | 15 |
| Kas-1 | Kas-1 | CS903 | NASC | 35 | 77 |
| Kas-2 | Kas-2 | CS76150 | NASC | 35 | 77 |
| Kolyv-6 | Kolyv-6 | CS76980 | NASC | 51.33 | 82.54 |
| Koren-1 | Koren-1 | CS76983 | NASC | 41.83 | 25.69 |
| Kz-13 | Kz-13 | CS76994 | NASC | 49.5 | 73.1 |
| Ler | Ler_Cybrid | CS2110199 | Laboratory of Genetics (WUR) | 47.984 | 10.8719 |
| Lesno-1 | Lesno-1 | CS77032 | NASC | 53.04 | 51.9 |
| Mammo-1 | Mammo-1 | CS76365 | NASC | 38.36 | 16.23 |
| Melni-2 | Melni-2 | CS77080 | NASC | 41.53 | 23.39 |
| Oua-0 | Oua-0 | CS799923 | NASC | 32.07853 | -6.275309 |
| Panke-1 | Panke-1 | CS77162 | NASC | 53.82 | 80.31 |
| Penbont-2 | - | - | John Doonan | - | - |
| Qar-8a | Qar-8a | CS76581 | NASC | 34.1 | 35.84 |
| Rab-2 | Madeira_13 | - | Maarten Koornneef | 32.754 | -17.13 |
| RRS-7 | RRS-7 | CS22564 | NASC | 41.5609 | -86.4251 |
| Samos-3a | - | - | Maarten Koornneef | 37.7992 | 26.8306 |
| Samos-4 | - | - | Maarten Koornneef | 37.7856 | 26.8467 |
| Sha | S-S_Cybrid | CS2110200 | Laboratory of Genetics (WUR) | 37.3 | 71.3 |
| Sij-2 | Sij-2 | CS76380 | NASC | 41.45 | 70.05 |
| Sij-4 | Sij-4 | CS9656 | NASC | 41.45 | 70.05 |
| Staro-2 | Staro-2 | CS77273 | NASC | 44.3 | 21.08 |
| Sus-1 | Sus-1 | CS76607 | NASC | 42.1833 | 73.4 |
| Tanz-1 | Tanz-1 | CS75719 | Laboratory of Genetics (WUR) | -2.8739 | 36.12 |
| Tanz-2 | - | CS75720 | Laboratory of Genetics (WUR) | -2.8739 | 36.12 |
| Taz-0 | Taz-0 | CS799913 | NASC | 34.09166 | -4.10258 |
| Toufl | - | CS76348 | NASC | 31.47 | -7.42 |
| Ws-4 | Ws-4_Cybrid | CS2110201 | Laboratory of Genetics (WUR) | 52.5 | 30 |
| Yeg-1 | Yeg-1 | CS75468 | NASC | 39.8692 | 45.3622 |
| Zin-9 | Zin-9 | CS799909 | NASC | 35.4528 | -5.42698 |

## Supplementary Figures

**Supplementary Figure 1.**
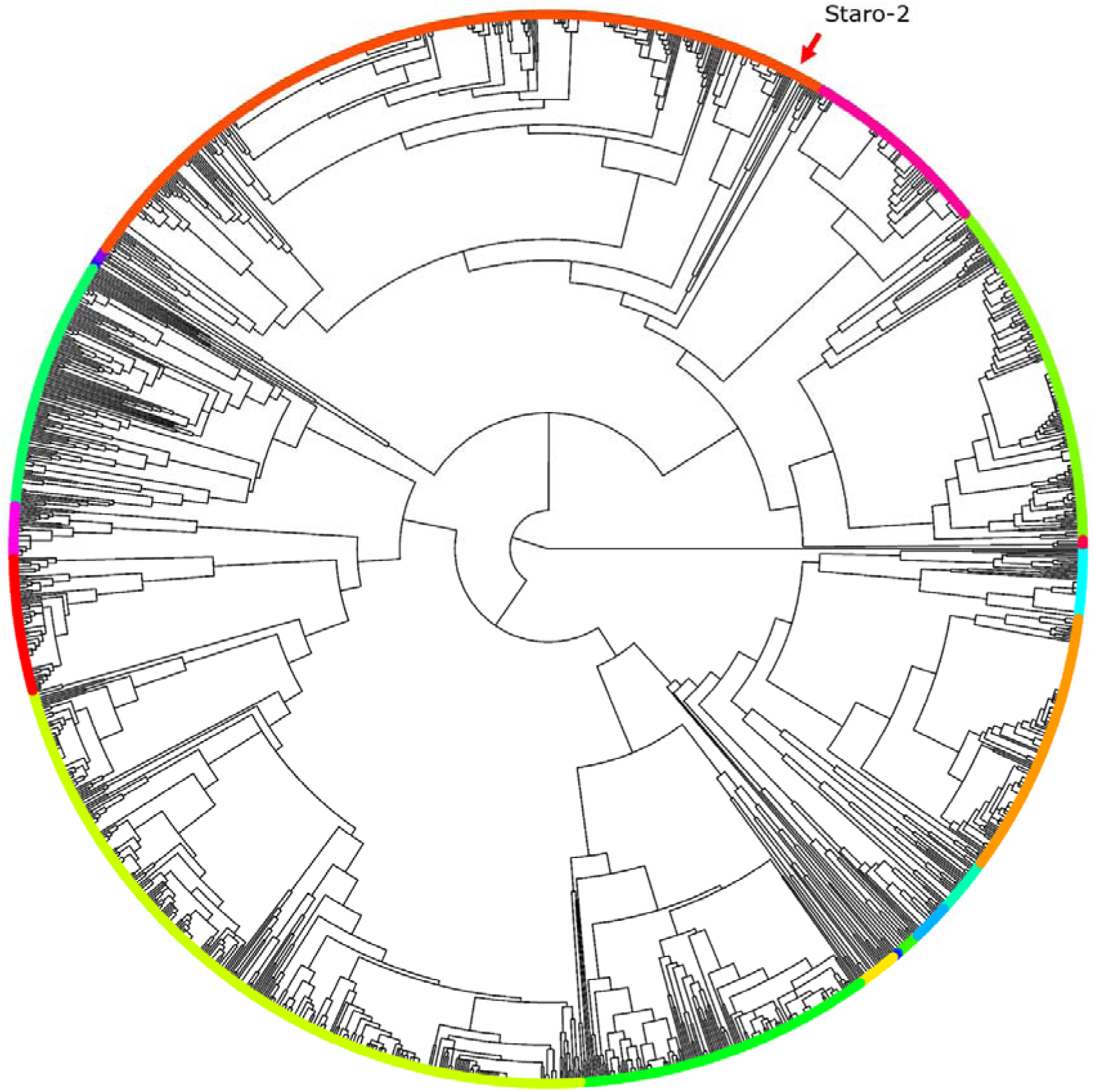
Neighbour joining tree for the chloroplast genome of 1531 accessions. The colours represent the subclusters (k=20) as in Figure 1B, but unrooted. The colours used for an accession match the colours used in Supplementary Figures 2 and 3. The red arrow indicates the position of Staro-2.

**Supplementary Figure 2.**
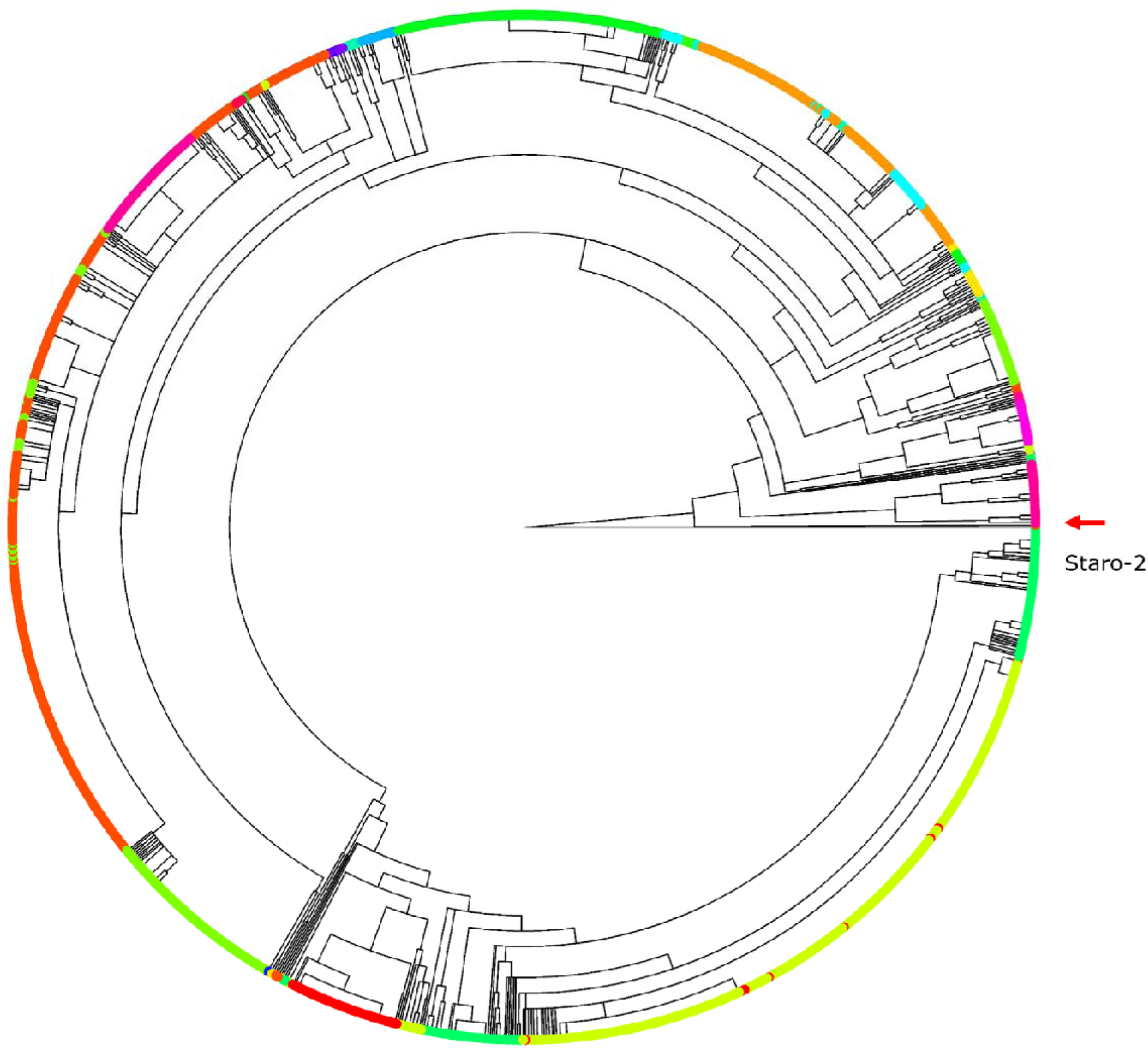
Neighbour joining tree for the mitochondrial genome of 1531 accessions. Tree displayed in fan form, and colour coded with the k=20 subclusters as identified based on the chloroplast analysis (Supplementary Figure 1). The overlap between the mitochondrial and chloroplast neighbour joining tree can be seen how the colours of the 20 clusters align with groups in the mitochondrial neighbour joining tree. The red arrow indicates the position of Staro-2.

**Supplementary Figure 3.**
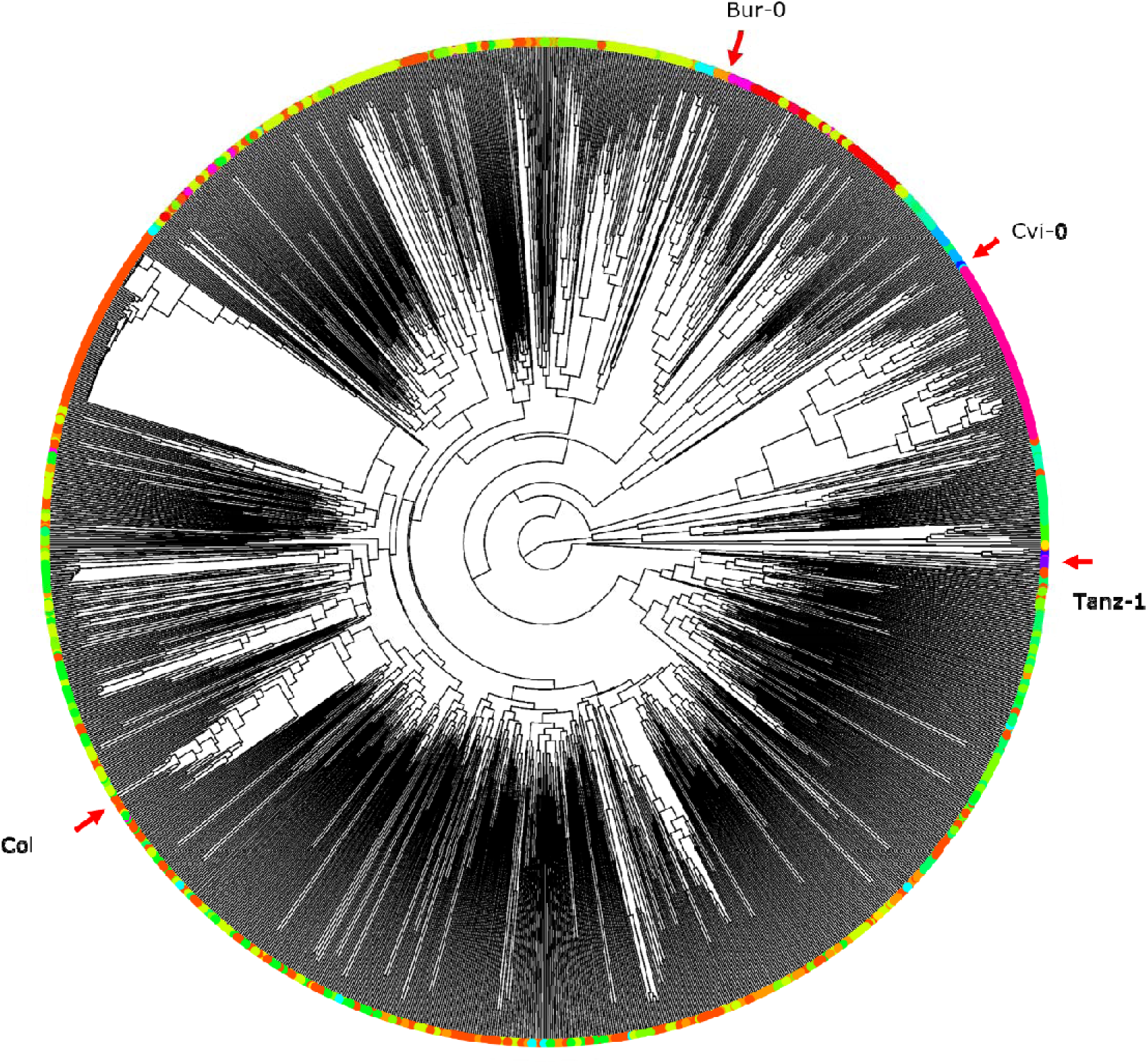
Neighbour joining tree for the nuclear genome of 1531 accessions. Tree displayed in fan form, and colour coded with the k=20 subclusters as identified based on the chloroplast genome analysis (Supplementary Figure 1). The arrows indicate the four nucleotypes used in the cybrid panel.

**Supplementary Figure 4.**
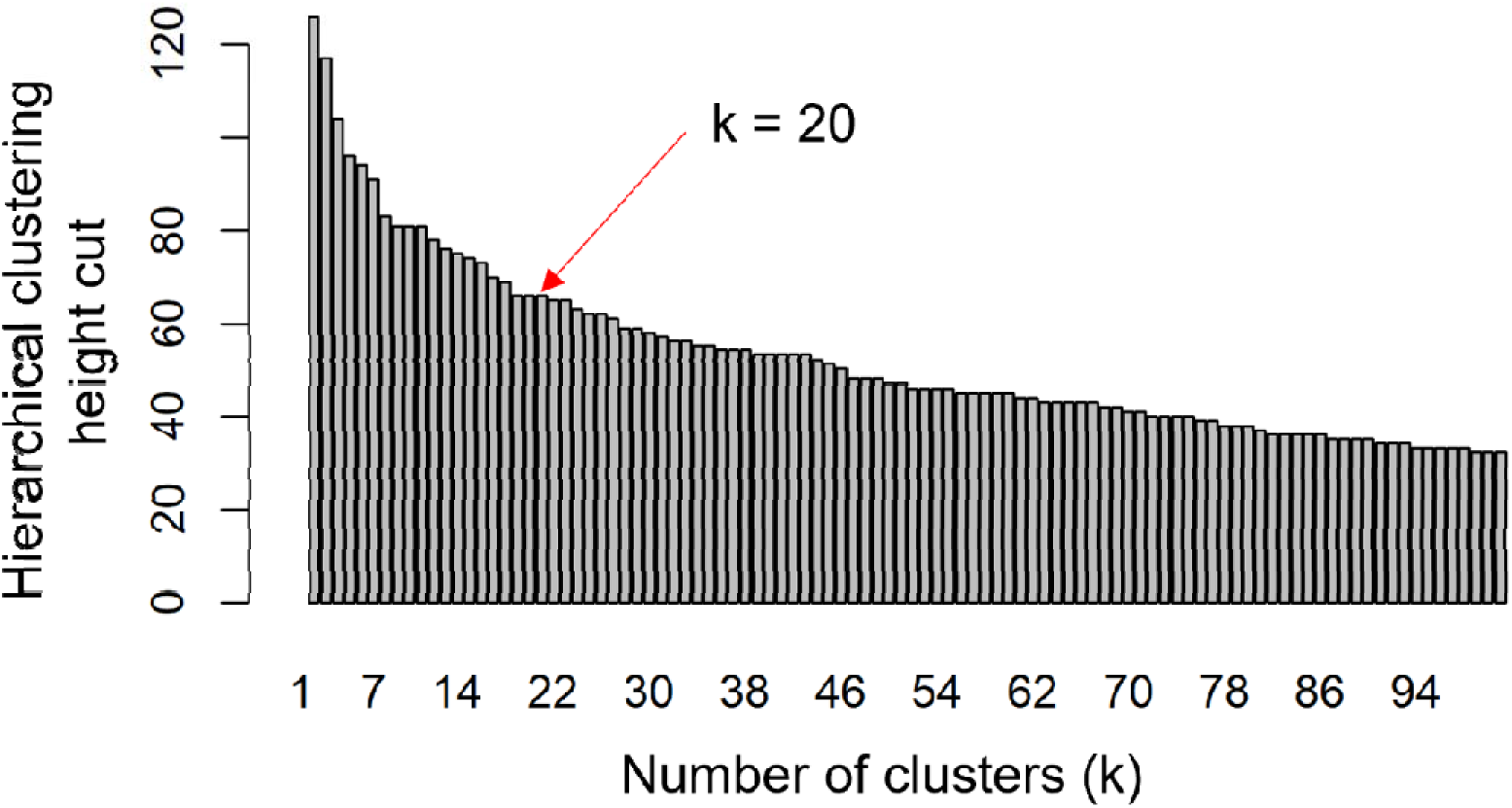
Elbow method to visualize the how the number of clusters influences the height of the cut-off. For this the hierarchical clustering data of the chloroplast genetic diversity is used. Based on this graph we arbitrarily chose k = 20.

**Supplementary Figure 5.**
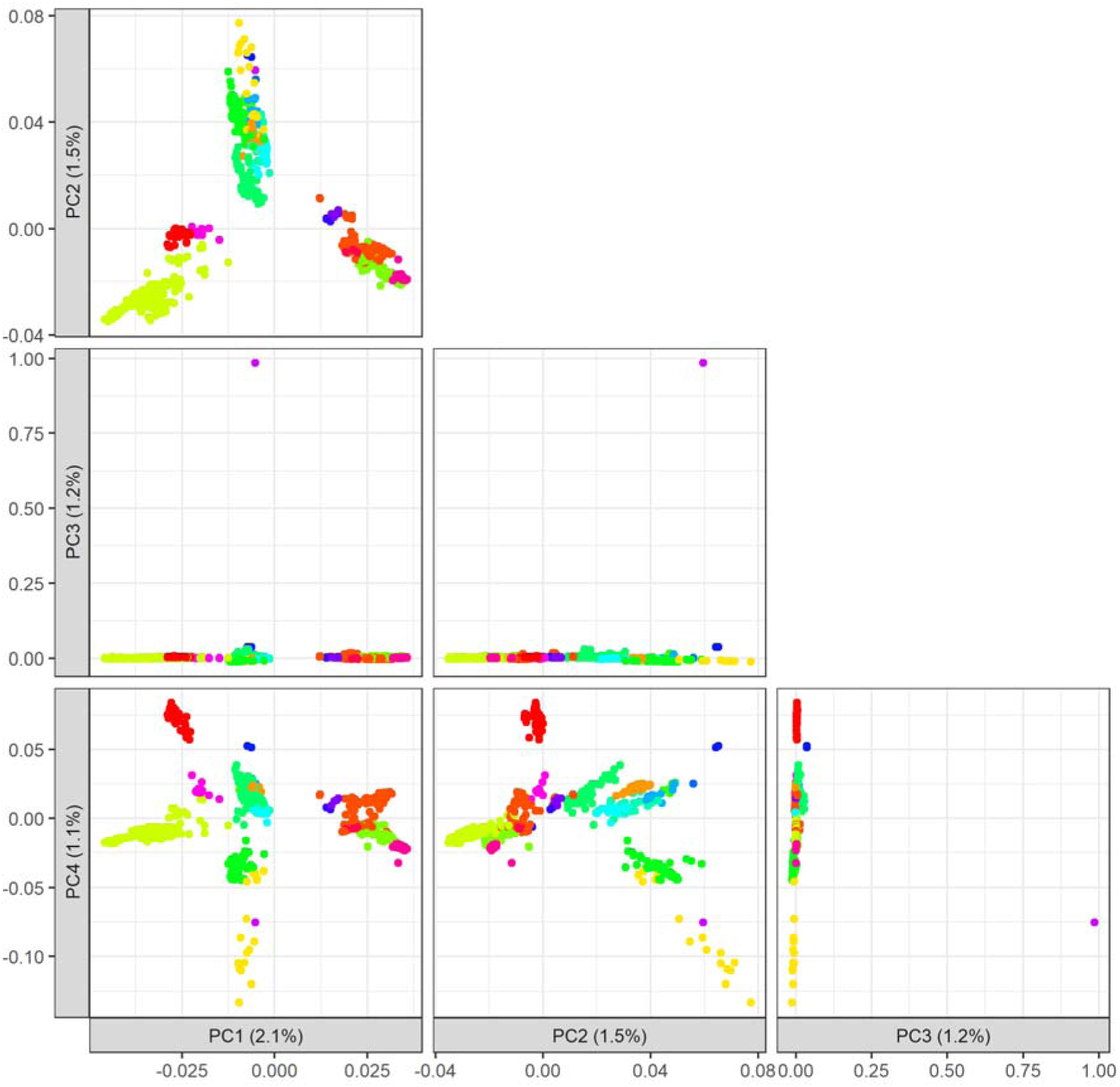
PC analysis based on the chloroplast genome of 1531 accessions. These PCs complement the first two PCs in Figure 1A. Colour codes are in line with the 20 subclusters as defined based on the chloroplast genome neighbour joining tree.

**Supplementary Figure 6.**
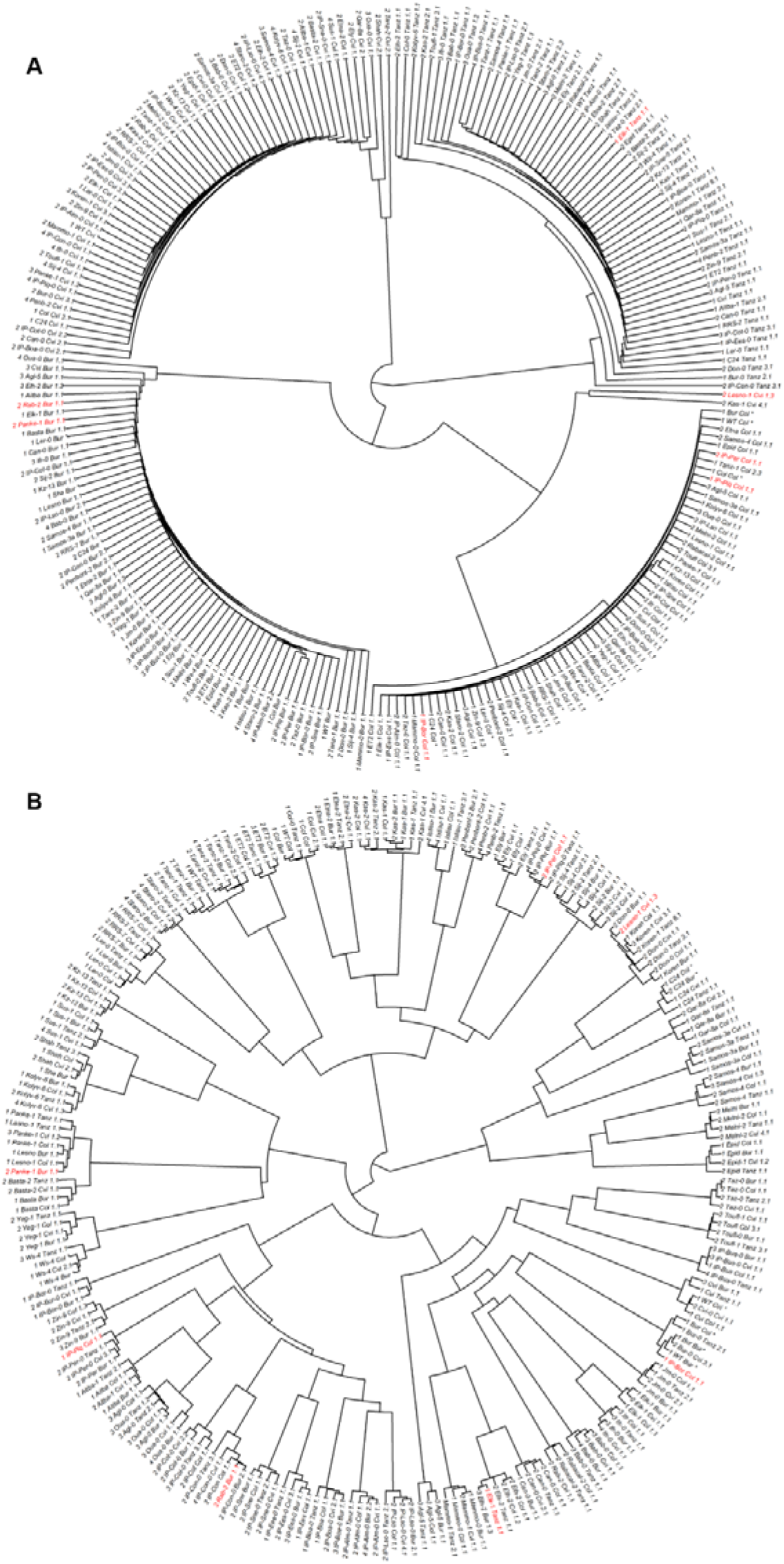
Genotyping of 240 novel cybrids via neighbour joining analysis of whole genomes sequencing data. A) Neighbour joining tree for the nuclear genetic variation, showing the four different nucleotypes. B) Neighbour joining tree for chloroplast variation, showing the separate clusters. Some clusters appear mixed, but this is due to relatively little or no genetic variation between given chloroplast genomes. In both plots the cybrids in red are excluded in further analyses.

**Supplementary Figure 7.**
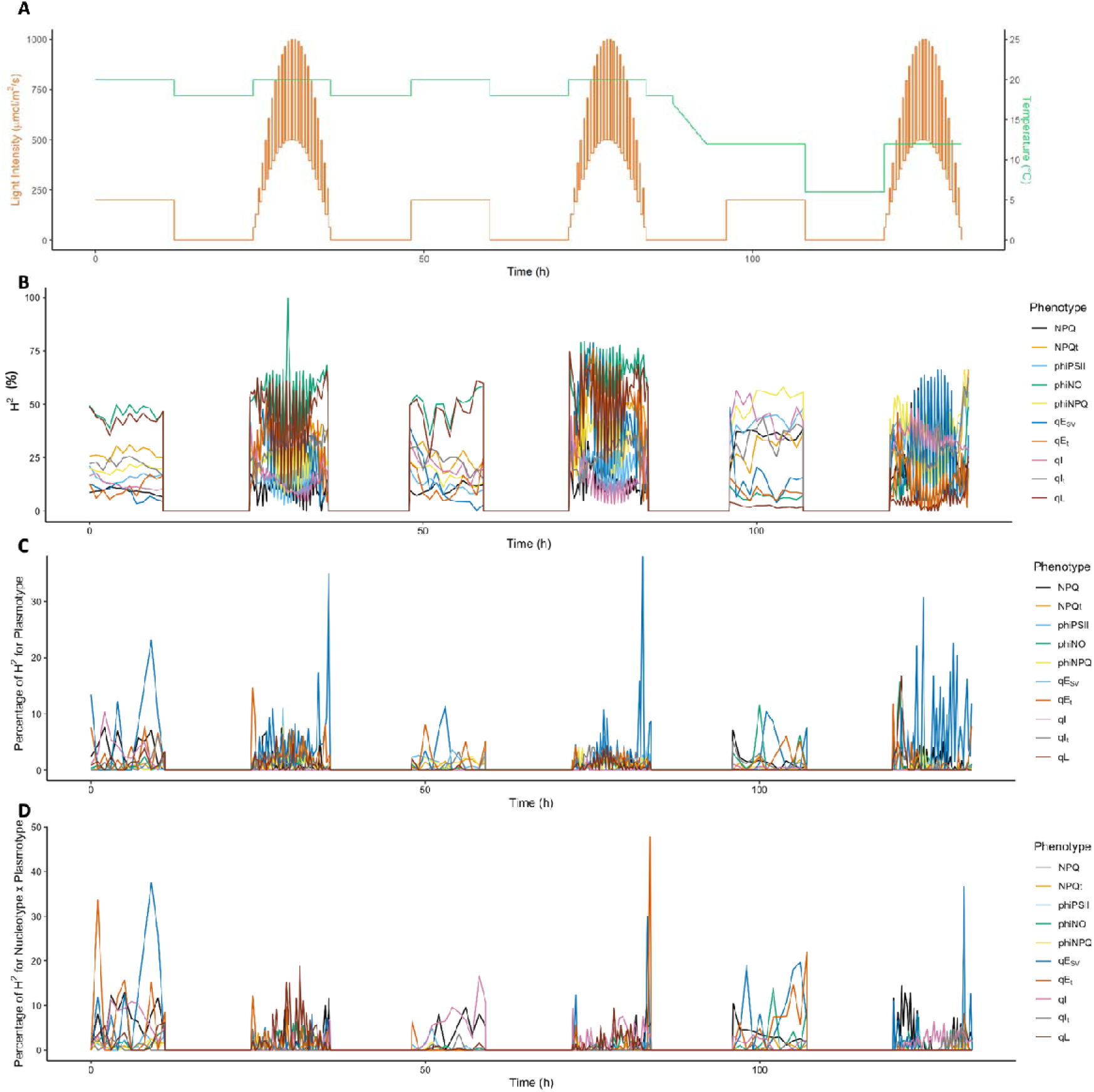
Overview of broad sense heritability, and the fraction explained by the plasmotype and nucleotype-plasmotype interaction components, in response to different environmental conditions in the DEPI system. A) Environmental conditions to which the plants were exposed after being grown under steady-state light conditions of 200 µmol m^-2^ s^-1^, with t=0 in this plot being the morning of day 21 after sowing. B) Broad sense heritability (H^2^; shown as percentage) for different photosynthetic parameters in response to the different environmental conditions, as shown in panel A. C) Percentage H^2^ for different photosynthetic parameters in response to the different environmental conditions for the additive plasmotype effect. D) The same as panel C, but for the percentage H^2^ for different photosynthetic parameters in response to the different environmental conditions for the nucleotype-plasmotype interaction effect.

**Supplementary Figure 8.**
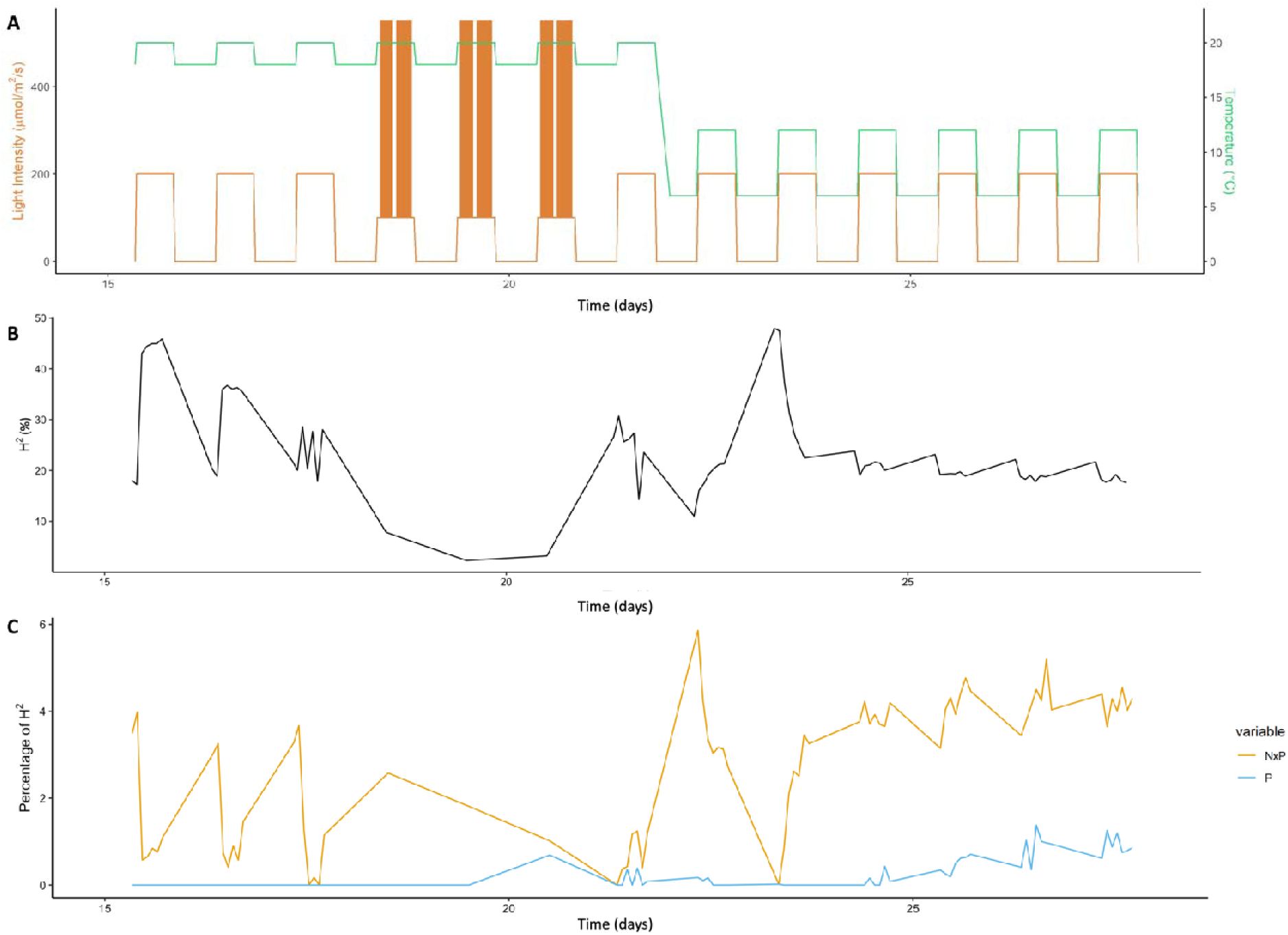
Overview of broad sense heritability, and the fraction explained by the plasmotype and nucleotype-plasmotype interaction components, in response to different environmental conditions in the Phenovator system. A) Environmental conditions to which the plants were exposed. The days before 15 days after sowing were identical to day 15. B) Broad sense heritability (H^2^; shown as percentage) for Φ_PSII_ in response to the different environmental conditions, as shown in panel A. C) Percentage H^2^ for Φ_PSII_ in response to the different environmental conditions for the additive plasmotype effect (blue) and nucleotype-plasmotype interaction effect (yellow).

**Supplementary Figure 9.**
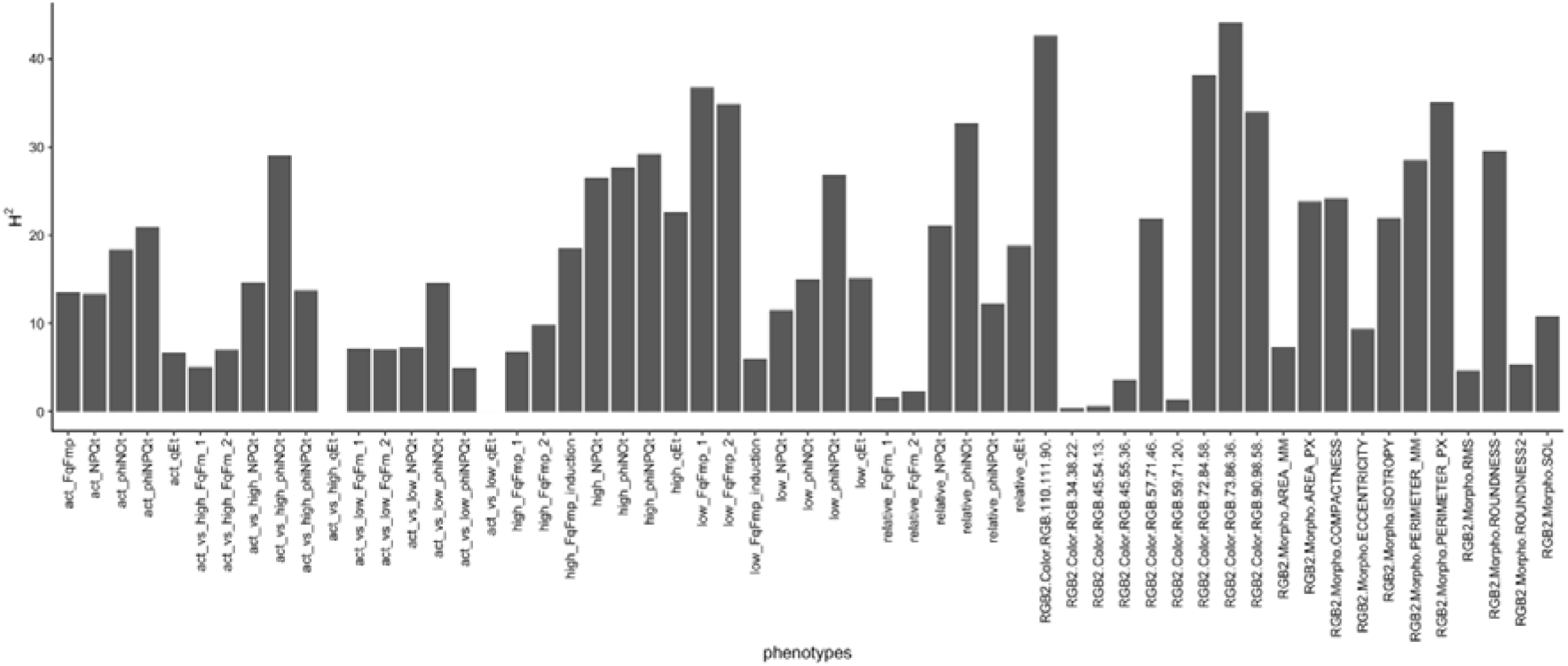
Overall H^2^ for phenotypes measured over two consecutive years in the semi-protected gauze tunnel. Morphological phenotypes start with “RGB”.

**Supplementary Figure 10.**
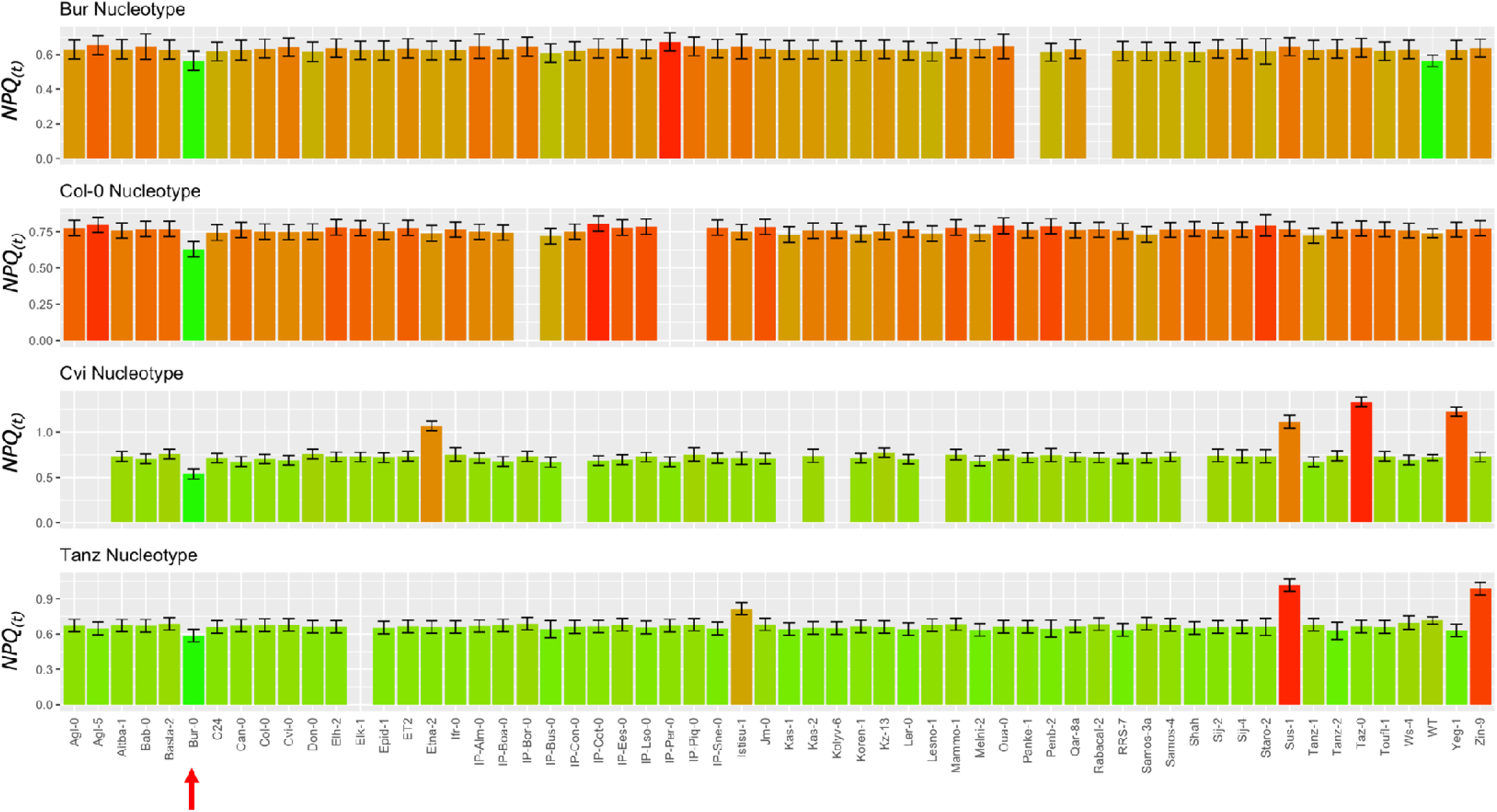
Bur-0 plasmotype (red arrow) effect as observed in the semi-natural outside experiment (combined over the two years). While visible in many timepoints, NPQ_(t)_ during low light after a high light period showed the biggest effect size difference. Note also how the wildtype plasmotype, i.e. Bur-0, in the Bur-0 nucleotype responds the same as the Bur-0^Bur-0^ self-cybrid. The same plasmotypes, although in different nucleotypes, are placed above each other. The cybrids are divided in four rows of bar graphs, with each row representing one nucleotype (which nucleotype is stated above). The bars are coloured per nucleotype on a scale with the lowest value as green and highest value as red. The error bars represent the standard error of the mean.

**Supplementary Figure 11.**
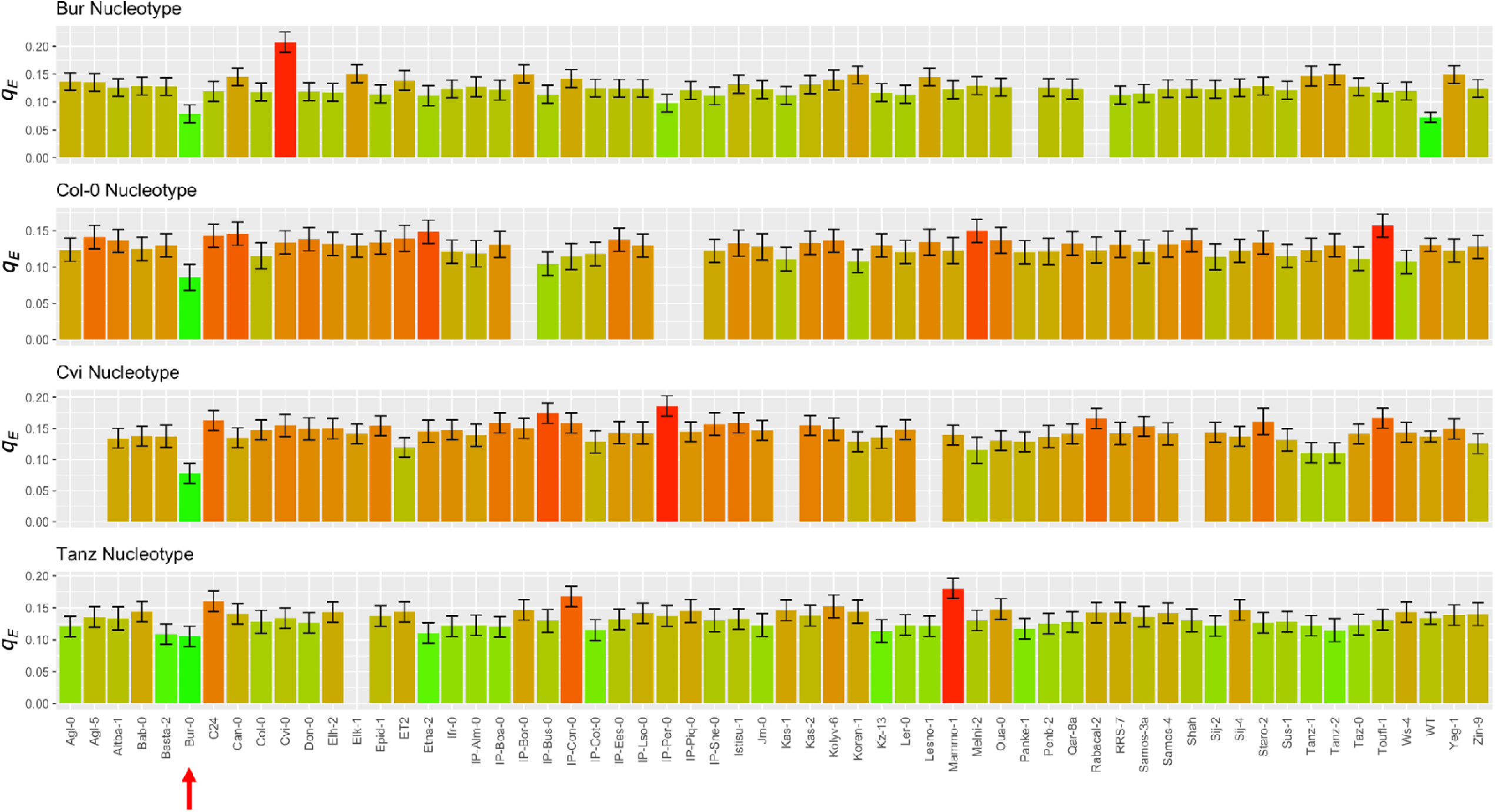
Bur-0 plasmotype (red arrow) effect during fluctuating light conditions of DEPI for q_E_. The same plasmotypes, although in different nucleotypes, are placed above each other. The cybrids are divided in four rows of bar graphs, with each row representing one nucleotype (which nucleotype is stated above). The bars are coloured per nucleotype on a scale with the lowest value as green and highest value as red. The error bars represent the standard error of the mean.

**Supplementary Figure 12.**
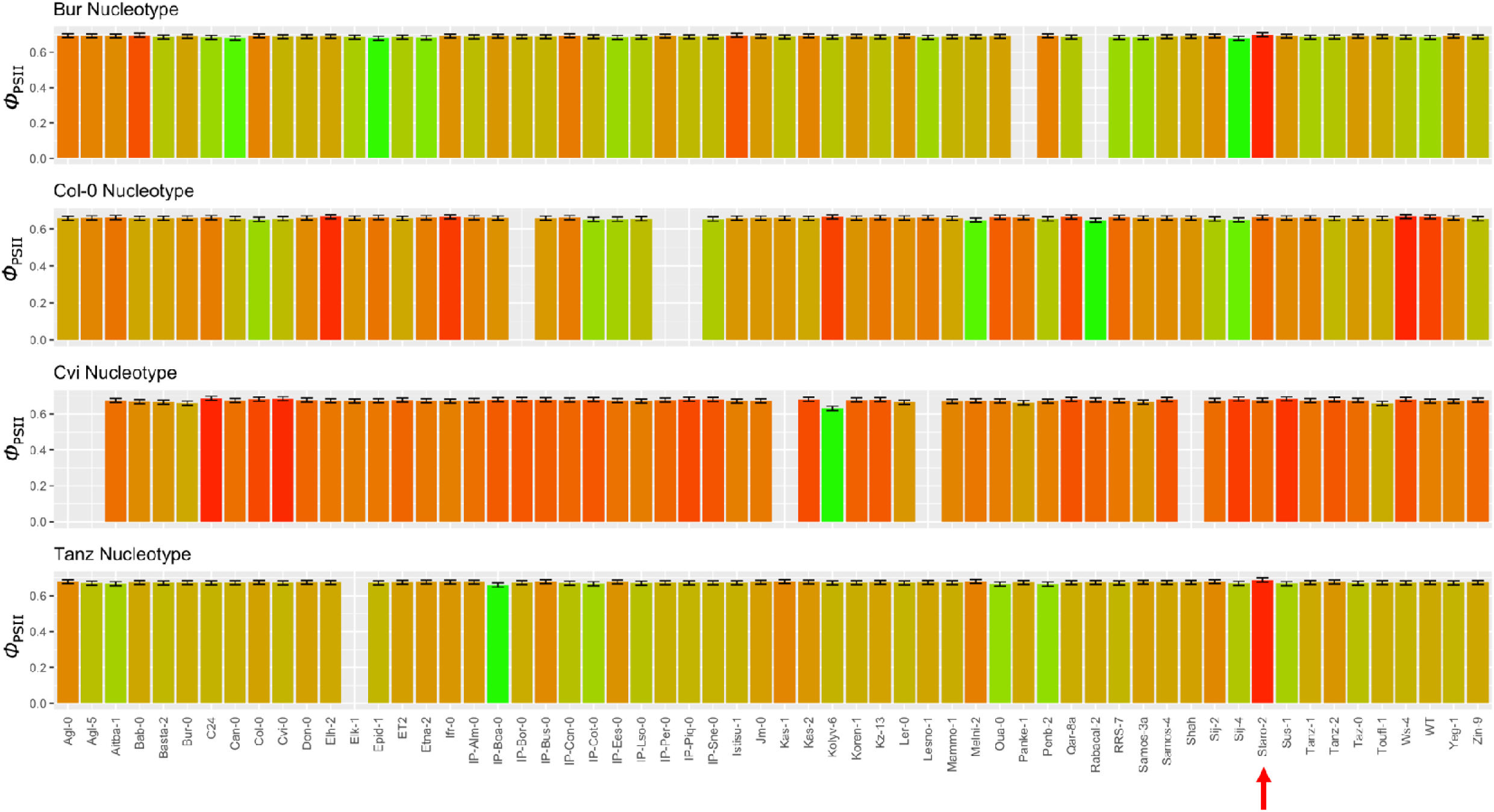
Staro-2 plasmotype (red arrow) effect on Φ_PSII_ measured on day 16 after sowing in the Phenovator experiment (grown in stable light at 200 µmol m^-2^ s^-1^). The same plasmotypes, although in different nucleotypes, are place above each other. The cybrids are divided in four rows of bar graphs, with each row representing one nucleotype (which nucleotype is stated above). The bars are coloured per nucleotype on a scale with the lowest value as green and highest value as red. The error bars represent the standard error of the mean.

**Supplementary Figure 13.**
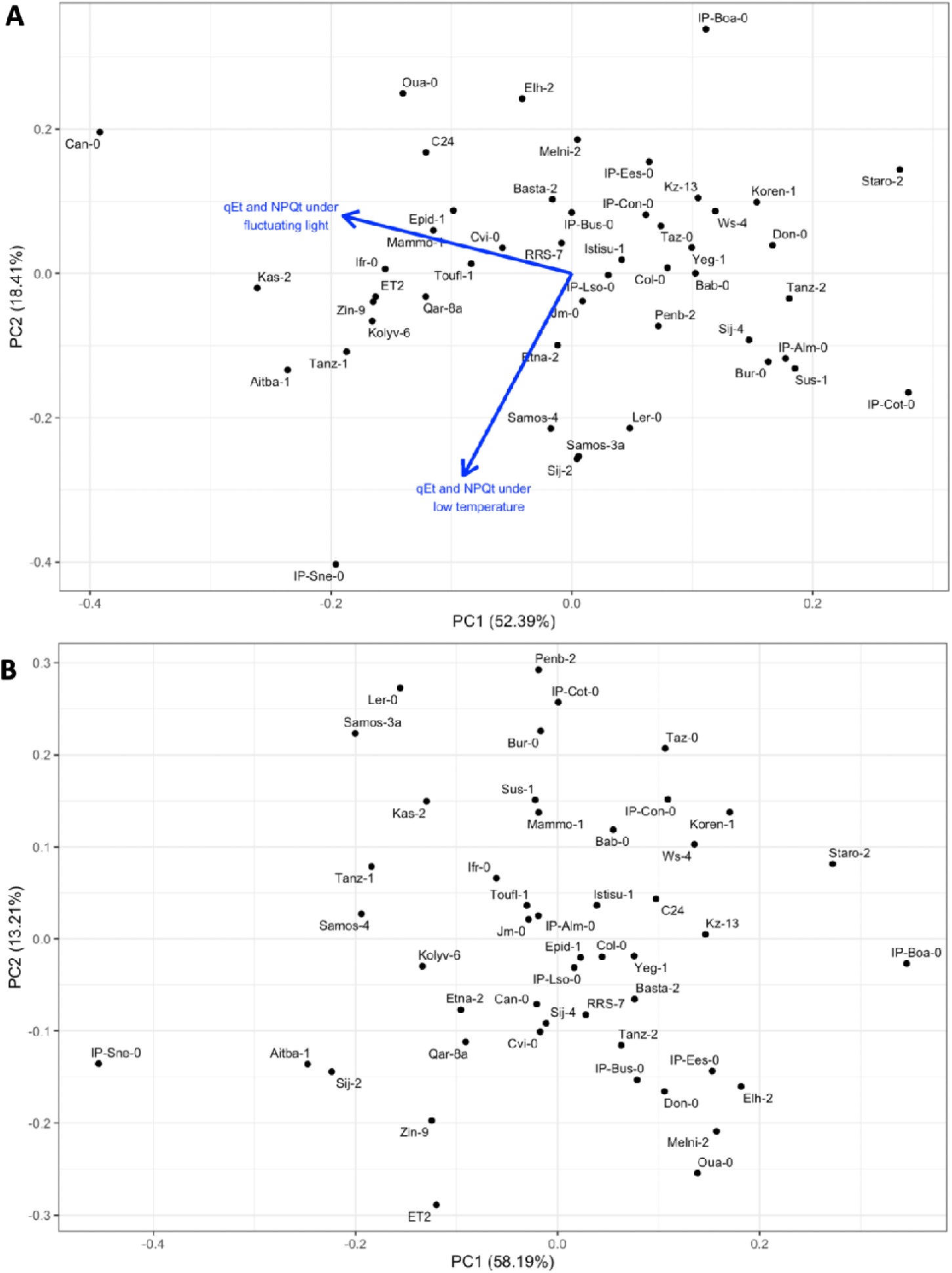
The plasmotype effect induced by IP-Sne and IP-Boa. A) The PC plot for the photosynthetic phenotypes of the DEPI experiment (as shown in Figure 4A), with the two main eigenvectors plotted onto it. B) The PC plot for the photosynthetic phenotypes during the fluctuating light conditions in cold temperatures.

**Supplementary Figure 14.**
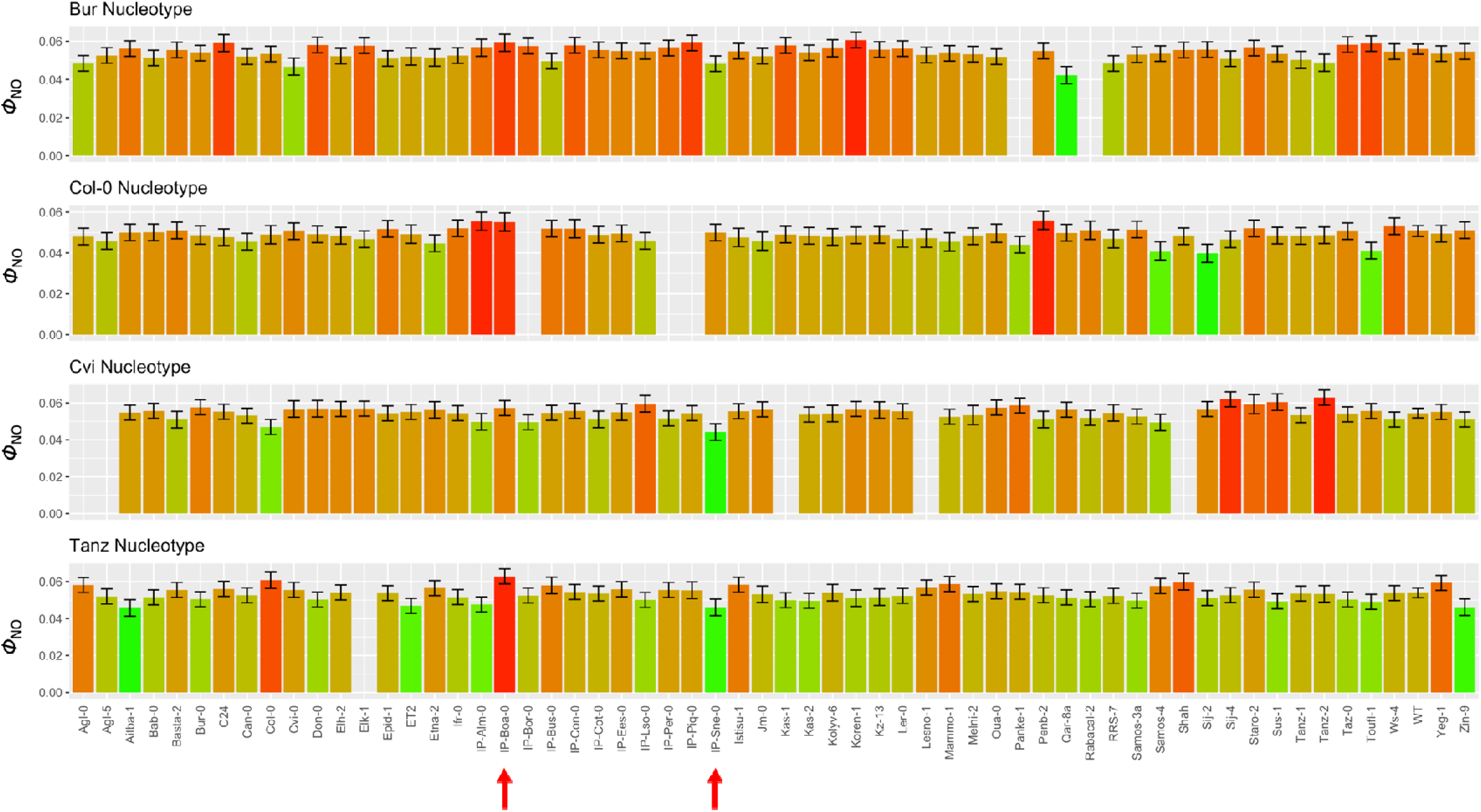
The response of Φ_NO_ at 128.716 h during the fluctuating light conditions in cold temperatures, with the arrows indicating the IP-Sne and IP-Boa plasmotypes. The cybrids are divided in four rows of bar graphs, with each row representing one nucleotype (which nucleotype is stated above). The bars are coloured per nucleotype on a scale with the lowest value as green and highest value as red. The error bars represent the standard error of the mean.

**Supplementary Figure 15.**
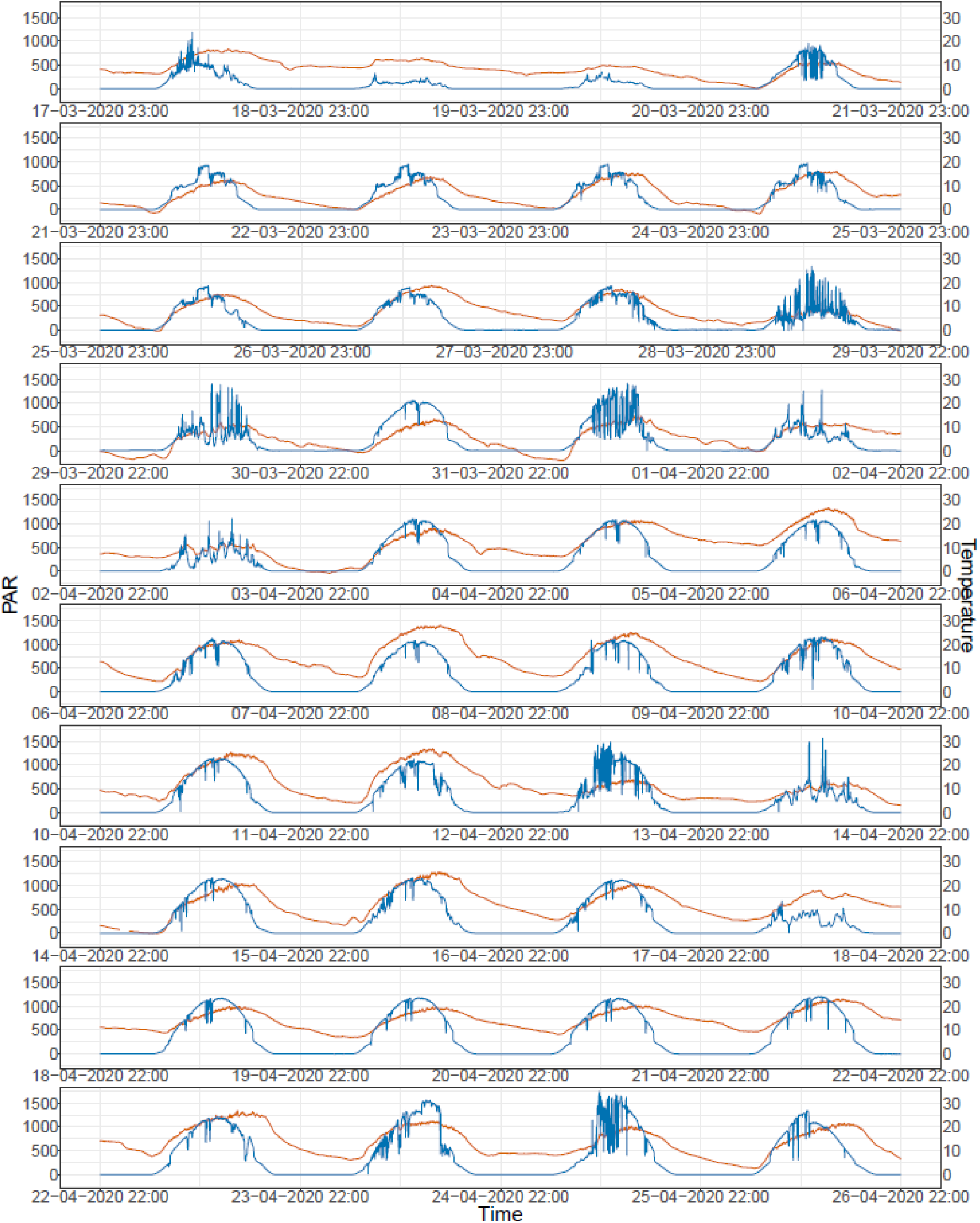
Light intensity (µmol m^-2^s^-1^) and temperature for the semi-protected experiment in spring 2020. Ligh intensity in blue and temperature in orange.

**Supplementary Figure 16.**
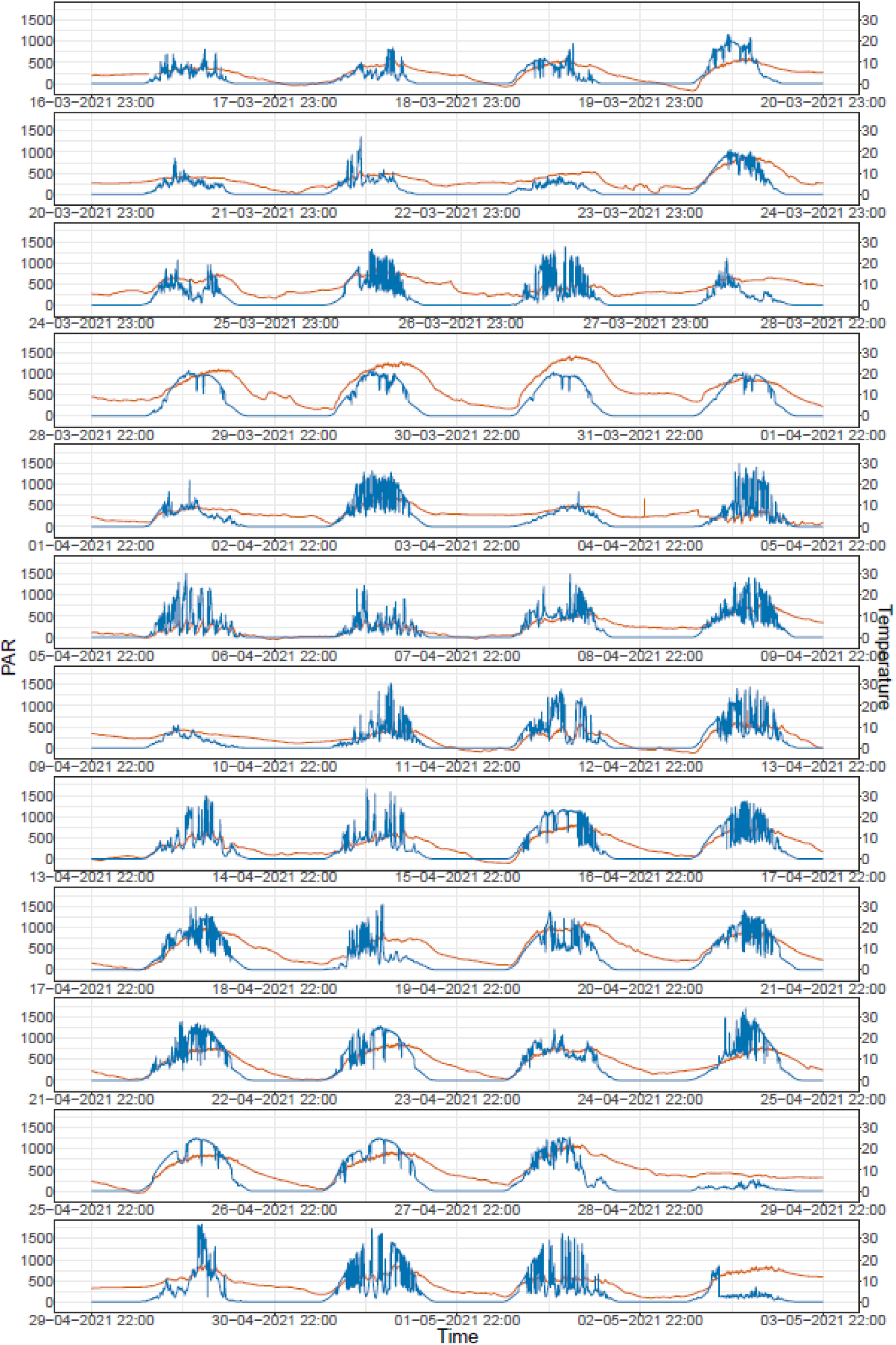
Light intensity (µmol m^-2^s^-1^) and temperature for the semi-protected experiment in spring 2021. Light intensity in blue and temperature in orange.

## References

1. S. P. Long, X. G. Zhu, S. L. Naidu, D. R. Ort, Can improvement in photosynthesis increase crop yields? Plant, Cell and Environment 29, 315–330 (2006).

2. X. G. Zhu, S. P. Long, D. R. Ort, Improving photosynthetic efficiency for greater yield. Annual Review of Plant Biology 61, 235–261 (2010).

3. D. R. Ort, et al., Redesigning photosynthesis to sustainably meet global food and bioenergy demand. Proceedings of the National Academy of Sciences of the United States of America 112, 8529–8536 (2015).

4. A. Hitchcock, et al., Redesigning the photosynthetic light reactions to enhance photosynthesis – the PhotoRedesign consortium. Plant Journal 109, 23–34 (2022).

5. P. J. Flood, J. Harbinson, M. G. M. Aarts, Natural genetic variation in plant photosynthesis. Trends in Plant Science 16, 327–335 (2011).

6. T. P. J. M. Theeuwen, L. L. Logie, J. Harbinson, M. G. M. Aarts, Genetics as a key to improving crop photosynthesis. Journal Of Experimental Botany 73, 3122–3137 (2022).

7. S. M. Driever, T. Lawson, P. J. Andralojc, C. A. Raines, M. A. J. Parry, Natural variation in photosynthetic capacity, growth, and yield in 64 field-grown wheat genotypes. Journal of Experimental Botany 65, 4959–4973 (2014).

8. Q. Wang, et al., Genetic architecture of natural variation in rice nonphotochemical quenching capacity revealed by genome-wide association study. Frontiers in Plant Science 8, 1773 (2017).

9. R. Van Rooijen, et al., Natural variation of YELLOW SEEDLING1 affects photosynthetic acclimation of Arabidopsis thaliana. Nature Communications 8, 1–9 (2017).

10. C. G. Oakley, et al., Genetic basis of photosynthetic responses to cold in two locally adapted populations of Arabidopsis thaliana. Journal of Experimental Botany 69, 699–709 (2018).

11. A. E. Prinzenberg, M. Víquez-Zamora, J. Harbinson, P. Lindhout, S. van Heusden, Chlorophyll fluorescence imaging reveals genetic variation and loci for a photosynthetic trait in diploid potato. Physiologia Plantarum 164, 163–175 (2018).

12. A. E. Prinzenberg, L. Campos-Dominguez, W. Kruijer, J. Harbinson, M. G. M. Aarts, Natural variation of photosynthetic efficiency in Arabidopsis thaliana accessions under low temperature conditions. Plant Cell and Environment 43, 2000–2013 (2020).

13. T. Rungrat, et al., A Genome-Wide Association Study of Non-Photochemical Quenching in response to local seasonal climates in Arabidopsis thaliana. Plant Direct 3, e00138 (2019).

14. M. Faralli, J. Matthews, T. Lawson, Exploiting natural variation and genetic manipulation of stomatal conductance for crop improvement. Current Opinion in Plant Biology 49, 1–7 (2019).

15. S. Adachi, et al., Genetic architecture of leaf photosynthesis in rice revealed by different types of reciprocal mapping populations. Journal of Experimental Botany 70, 5131–5144 (2019).

16. K. Taniyoshi, Y. Tanaka, T. Shiraiwa, Genetic variation in the photosynthetic induction response in rice (Oryza sativa L.). Plant Production Science 513–521 (2020). 10.1080/1343943X.2020.1777878.

17. L. G. Acevedo-Siaca, et al., Variation in photosynthetic induction between rice accessions and its potential for improving productivity. New Phytologist 227, 1097–1108 (2020).

18. L. G. Acevedo-Siaca, R. Coe, W. P. Quick, S. P. Long, Variation between rice accessions in photosynthetic induction in flag leaves and underlying mechanisms. Journal of Experimental Botany 72, 1282–1294 (2021).

19. J. N. Timmis, M. A. Ayliff, C. Y. Huang, W. Martin, Endosymbiotic gene transfer: organelle genomes forge eukaryotic chromosomes. Nature Reviews Genetics 5, 123–135 (2004).

20. P. Lamesch, et al., The Arabidopsis Information Resource (TAIR): improved gene annotation and new tools. Nucleic Acids Research 40, D1202–D1210 (2012).

21. D. B. Sloan, Z. Wu, J. Sharbrough, Correction of Persistent Errors in Arabidopsis Reference Mitochondrial Genomes. The Plant Cell 30, 525–527 (2018).

22. D. O. Daley, J. Whelan, Why genes persist in organelle genomes. Genome Biology 6, 110 (2005).

23. D. G. Bock, R. L. Andrew, L. H. Rieseberg, On the adaptive value of cytoplasmic genomes in plants. Molecular Ecology 23, 4899–911 (2014).

24. D. C. Wallace, A Mitochondrial Paradigm of Metabolic and Degenerative Diseases, Aging, and Cancer: A Dawn for Evolutionary Medicine. Annual Review of Genetics 39, 359 (2005).

25. S. Greiner, “Plastome mutants of higher plants” in Genomics of Chloroplasts and Mitochondria, (Springer Science & Business Media, 2012), pp. 237–266.

26. M. W. Nachman, “Deleterious mutations in animal mitochondrial DNA” in Mutation and Evolution, Contemporary Issues in Genetics and Evolution., R. C. Woodruff, J. N. Thompson, Eds. (Springer Netherlands, 1998), pp. 61–69.

27. D. M. Rand, L. M. Kann, “Mutation and selection at silent and replacement sites in the evolution of animal mitochondrial DNA” in Mutation and Evolution, Contemporary Issues in Genetics and Evolution., R. C. Woodruff, J. N. Thompson, Eds. (Springer Netherlands, 1998), pp. 393–407.

28. K. H. Wolfe, W. H. Li, P. M. Sharp, Rates of nucleotide substitution vary greatly among plant mitochondrial, chloroplast, and nuclear DNAs. Proceedings of the National Academy of Sciences of the United States of America 84, 9054–9058 (1987).

29. M. E. El-Lithy, et al., Altered photosynthetic performance of a natural Arabidopsis accession is associated with atrazine resistance. Journal of Experimental Botany 56, 1625–1634 (2005).

30. P. J. Flood, et al., Whole-Genome Hitchhiking on an Organelle Mutation. Current Biology 26, 1306–1311 (2016).

31. M. V. Kapralov, D. A. Filatov, Widespread positive selection in the photosynthetic Rubisco enzyme. BMC Evolutionary Biology 2007 7:1 7, 1–10 (2007).

32. P. H. Leinonen, D. L. Remington, O. Savolainen, Local Adaptation, Phenotypic Differentiation, and Hybrid Fitness in Diverged Natural Populations of Arabidopsis Lyrata. Evolution 65, 90–107 (2011).

33. M. Moison, et al., Cytoplasmic phylogeny and evidence of cyto-nuclear co-adaptation in Arabidopsis thaliana. Plant Journal 63, 728–738 (2010).

34. Z. Yin, et al., Expression quantitative trait loci analysis of two genes encoding Rubisco activase in soybean. Plant Physiology 152, 1625–1637 (2010).

35. A. Tyagi, et al., Genetic diversity and population structure of Arabidopsis thaliana along an altitudinal gradient. AoB Plants 10, plv145 (2015).

36. C.-W. Hsu, C.-Y. Lo, C.-R. Lee, On the postglacial spread of human commensal Arabidopsis thaliana: journey to the East. New Phytologist 222, 1447–1457 (2019).

37. L. A. Corey, D. F. Matzinger, C. C. Cockerham, Maternal and reciprocal effects on seedling characters in Arabidopsis thaliana (L.) Heynh. Genetics 82, 677–683 (1976).

38. R. C. Meyer, O. Torjek, M. Becher, T. Altmann, Heterosis of biomass production in Arabidopsis. Establishment during early development. Plant Physiology 134, 1813–1823 (2004).

39. R. Fujimoto, J. M. Taylor, S. Shirasawa, W. J. Peacock, E. S. Dennis, Heterosis of Arabidopsis hybrids between C24 and Col is associated with increased photosynthesis capacity. Proceedings of the National Academy of Sciences 109, 7109–7114 (2012).

40. S. Atwell, et al., Genome-wide association study of 107 phenotypes in Arabidopsis thaliana inbred lines. Nature 465, 627–31 (2010).

41. V. S. Gordon, J. E. Staub, Comparative Analysis of Chilling Response in Cucumber Through Plastidic and Nuclear Genetic Effects Component Analysis. Journal of the American Society for Horticultural Science 136, 256–264 (2011).

42. M. Miclaus, et al., Maize Cytolines Unmask Key Nuclear Genes That Are under the Control of Retrograde Signaling Pathways in Plants. Genome Biology and Evolution 8, 3256–3270 (2016).

43. P. J. Flood, et al., Reciprocal cybrids reveal how organellar genomes affect plant phenotypes. Nature Plants 6, 13–21 (2020).

44. M. Ravi, S. W. L. Chan, Haploid plants produced by centromere-mediated genome elimination. Nature 464, 615–618 (2010).

45. C. Alonso-Blanco, et al., 1,135 Genomes Reveal the Global Pattern of Polymorphism in Arabidopsis thaliana. Cell 166, 481–491 (2016).

46. A. Durvasula, et al., African genomes illuminate the early history and transition to selfing in Arabidopsis thaliana. Proceedings of the National Academy of Sciences 201616736 (2017). 10.1073/PNAS.1616736114.

47. L. Zeng, et al., Discovery of a high-altitude ecotype and ancient lineage of Arabidopsis thaliana from Tibet. Science Bulletin 62, 1628–1630 (2017).

48. Y. P. Zou, et al., Adaptation of Arabidopsis thaliana to the Yangtze River basin. Genome Biology 18, 1–11 (2017).

49. A. Fulgione, M. Koornneef, F. Roux, J. Hermisson, A. M. Hancock, Madeiran Arabidopsis thaliana Reveals Ancient Long-Range Colonization and Clarifies Demography in Eurasia. Molecular Biology and Evolution 35, 564–574 (2018).

50. J.-H. Xu, et al., Dynamics of chloroplast genomes in green plants. Genomics 106, 221–231 (2015).

51. J. M. Gualberto, K. J. Newton, Plant Mitochondrial Genomes: Dynamics and Mechanisms of Mutation. Annual Review of Plant Biology 68, 225–252 (2017).

52. Y. Zou, W. Zhu, D. B. Sloan, Z. Wu, Long-read sequencing characterizes mitochondrial and plastid genome variants in Arabidopsis msh1 mutants. The Plant Journal 112, 738–755 (2022).

53. V. Shedge, M. Arrieta-Montiel, A. C. Christensen, S. A. Mackenzie, Plant Mitochondrial Recombination Surveillance Requires Unusual RecA and MutS Homologs. The Plant Cell 19, 1251–1264 (2007).

54. M. P. Arrieta-Montiel, V. Shedge, J. Davila, A. C. Christensen, S. A. Mackenzie, Diversity of the Arabidopsis Mitochondrial Genome Occurs via Nuclear-Controlled Recombination Activity. Genetics 183, 1261–1268 (2009).

55. M. Ravi, S. W. L. Chan, Haploid plants produced by centromere-mediated genome elimination. Nature 464, 615–618 (2010).

56. J. A. Cruz, et al., Dynamic Environmental Photosynthetic Imaging Reveals Emergent Phenotypes. Cell Systems 2, 365–377 (2016).

57. P. J. Flood, et al., Phenomics for photosynthesis, growth and reflectance in Arabidopsis thaliana reveals circadian and long-term fluctuations in heritability. Plant Methods 12, 14 (2016).

58. T. P. J. M. Theeuwen, et al., The NDH complex reveals a trade-off preventing maximizing photosynthesis in Arabidopsis thaliana. [Preprint] (2022).

59. D. M. Kramer, G. Johnson, O. Kiirats, G. E. Edwards, New Fluorescence Parameters for the Determination of Q _A_ Redox State and Excitation Energy Fluxes. Photosynthesis Research 79, 209–218 (2004).

60. J. Provan, J. J. Campanella, Patterns of Cytoplasmic Variation in Arabidopsis thaliana (Brassicaceae) Revealed by Polymorphic Chloroplast Microsatellites. Systematic Botany 28, 578–583 (2003).

61. F. X. Picó, B. Méndez-Vigo, J. M. Martínez-Zapater, C. Alonso-Blanco, Natural Genetic Variation of Arabidopsis thaliana Is Geographically Structured in the Iberian Peninsula. Genetics 180, 1009–1021 (2008).

62. A. K. Azhagiri, P. Maliga, Exceptional paternal inheritance of plastids in Arabidopsis suggests that low-frequency leakage of plastids via pollen may be universal in plants. The Plant Journal 52, 817–823 (2007).

63. K. H. Wolfe, W. H. Li, P. M. Sharp, Rates of nucleotide substitution vary greatly among plant mitochondrial, chloroplast, and nuclear DNAs. Proceedings of the National Academy of Sciences of the United States of America 84, 9054–9058 (1987).

64. G. Drouin, H. Daoud, J. Xia, Relative rates of synonymous substitutions in the mitochondrial, chloroplast and nuclear genomes of seed plants. Molecular Phylogenetics and Evolution 49, 827–831 (2008).

65. C.-R. Lee, et al., On the post-glacial spread of human commensal Arabidopsis thaliana. Nature Communications 8, 14458 (2017).

66. F. Roux, et al., Cytonuclear interactions affect adaptive traits of the annual plant Arabidopsis thaliana in the field. Proceedings of the National Academy of Sciences of the United States of America 113, 3687–92 (2016).

67. Y. Fan, et al., The crucial roles of mitochondria in supporting C4 photosynthesis. New Phytologist 233, 1083–1096 (2021).

68. T. P. J. M. Theeuwen, Exploring the unexplored: Unravelling the cyto-nuclear interactions in Arabidopsis thaliana to improve photosynthesis. (2023). 10.18174/583113.

69. L. Chen, Y.-G. Liu, Male Sterility and Fertility Restoration in Crops. Annual Review of Plant Biology 65, 579–606 (2014).

70. R. Horn, K. J. Gupta, N. Colombo, Mitochondrion role in molecular basis of cytoplasmic male sterility. Mitochondrion 19, 198–205 (2014).

71. M. Sabar, R. De Paepe, Y. de Kouchkovsky, Complex I Impairment, Respiratory Compensations, and Photosynthetic Decrease in Nuclear and Mitochondrial Male Sterile Mutants of *Nicotiana sylvestris*. Plant Physiology 124, 1239–1250 (2000).

72. E. H. Murchie, A. V. Ruban, Dynamic non-photochemical quenching in plants: from molecular mechanism to productivity. The Plant Journal 101, 885–896 (2020).

73. D. D. Strand, L. D’Andrea, R. Bock, The plastid NAD(P)H dehydrogenase-like complex: structure, function and evolutionary dynamics. Biochemical Journal 476, 2743–2756 (2019).

74. A. Korte, A. Farlow, The advantages and limitations of trait analysis with GWAS: a review. Plant Methods 2013 9:1 9, 1–9 (2013).

75. P. E. Chen, B. J. Shapiro, The advent of genome-wide association studies for bacteria. Current Opinion in Microbiology 25, 17–24 (2015).

76. M. M. Maury, et al., Uncovering Listeria monocytogenes hypervirulence by harnessing its biodiversity. Nature Genetics 48, 308–313 (2016).

77. F. Coll, et al., PowerBacGWAS: a computational pipeline to perform power calculations for bacterial genome-wide association studies. Communications Biology 5, 1–12 (2022).

78. S. Ruf, et al., High-efficiency generation of fertile transplastomic Arabidopsis plants. Nature Plants 5, 282–289 (2019).

79. S. Y. Kwak, et al., Chloroplast-selective gene delivery and expression in planta using chitosan-complexed single-walled carbon nanotube carriers. Nature Nanotechnology 14, 447–455 (2019).

80. A. Jakubiec, et al., Replicating minichromosomes as a new tool for plastid genome engineering. Nature Plants 7, 932–941 (2021).

81. B. C. Kang, et al., Chloroplast and mitochondrial DNA editing in plants. Nature Plants 7, 899–905 (2021).

82. I. Nakazato, et al., Targeted base editing in the plastid genome of Arabidopsis thaliana. Nature Plants 7, 906–913 (2021).

83. F. Budar, F. Roux, The role of organelle genomes in plant adaptation. Plant Signaling & Behavior 6, 635–639 (2011).

84. J. Lv, et al., Generation of paternal haploids in wheat by genome editing of the centromeric histone CENH3. Nature Biotechnology 2020 38:12 38, 1397–1401 (2020).

85. N. Wang, J. I. Gent, R. K. Dawe, Haploid induction by a maize cenh3 null mutant. Science Advances 7, eabe2299 (2021).

86. E. Bortiri, et al., Cyto-swapping in maize by haploid induction with a cenh3 mutant. Nat. Plants 10, 567–571 (2024).

87. F. Han, et al., One-step creation of CMS lines using a BoCENH3-based haploid induction system in Brassica crop. Nat. Plants 10, 581–586 (2024).

88. R. Y. Wijfjes, Computational analysis of copy number variation in plant genomes. (2021). 10.18174/553649.

89. E. Paradis, K. Schliep, ape 5.0: an environment for modern phylogenetics and evolutionary analyses in R. Bioinformatics 35, 526–528 (2019).

90. B. J. Knaus, N. J. Grünwald, vcfr: a package to manipulate and visualize variant call format data in R. Molecular Ecology Resources 17, 44–53 (2017).

91. X. Zheng, et al., SeqArray—a storage-efficient high-performance data format for WGS variant calls. Bioinformatics 33, 2251–2257 (2017).

92. X. Zheng, et al., A high-performance computing toolset for relatedness and principal component analysis of SNP data. Bioinformatics 28, 3326–3328 (2012).

93. D. Gaio, et al., Hackflex: low-cost, high-throughput, Illumina Nextera Flex library construction. Microbial Genomics 8, 000744 (2022).

94. M. Martin, Cutadapt removes adapter sequences from high-throughput sequencing reads. EMBnet.journal 17, 10–12 (2011).

95. H. Li, Aligning sequence reads, clone sequences and assembly contigs with BWA-MEM. (2013). 10.48550/arxiv.1303.3997.

96. C. Chiang, et al., SpeedSeq: ultra-fast personal genome analysis and interpretation. Nature Methods 12, 966–968 (2015).

97. H. Li, et al., The Sequence Alignment/Map format and SAMtools. Bioinformatics 25, 2078–2079 (2009).

98. G. A. Van der Auwera, et al., From FastQ data to high confidence variant calls: the Genome Analysis Toolkit best practices pipeline. Current Protocols in Bioinformatics 11, 11.10.1–11.10.33 (2013).

99. E. Garrison, G. Marth, Haplotype-based variant detection from short-read sequencing. [Preprint] (2012). Available at: http://arxiv.org/abs/1207.3907.

100. C. C. Chang, et al., Second-generation PLINK: rising to the challenge of larger and richer datasets. GigaScience 4, s13742–015-0047–8 (2015).

101. E. Paradis, K. Schliep, ape 5.0: an environment for modern phylogenetics and evolutionary analyses in R. Bioinformatics 35, 526–528 (2019).

102. A. Junker, et al., Optimizing experimental procedures for quantitative evaluation of crop plant performance in high throughput phenotyping systems. Frontiers in Plant Science 5 (2015).

103. D. Bates, M. Mächler, B. Bolker, S. Walker, Fitting Linear Mixed-Effects Models using lme4. [Preprint] (2014). Available at: http://arxiv.org/abs/1406.5823 [Accessed 7 November 2022].

104. S. Tietz, C. C. Hall, J. A. Cruz, D. M. Kramer, NPQt: a chlorophyll fluorescence parameter for rapid estimation and imaging of non-photochemical quenching of excitons in photosystem-II-associated antenna complexes. Plant, Cell & Environment 40, 1243–1255 (2017).

105. X. Zhou, M. Stephens, Genome-wide efficient mixed-model analysis for association studies. Nat Genet 44, 821–824 (2012).

